# *Plasmodium falciparum* guanylyl cyclase-alpha and the activity of its appended P4-ATPase domain are essential for cGMP synthesis and blood stage egress

**DOI:** 10.1101/2020.09.07.285734

**Authors:** Stephanie D. Nofal, Avnish Patel, Michael J. Blackman, Christian Flueck, David A. Baker

**Affiliations:** Faculty of Infectious and Tropical Diseases, London School of Hygiene & Tropical Medicine, London, United Kingdom.; Malaria Biochemistry Laboratory, The Francis Crick Institute, London, United Kingdom.

## Abstract

In malaria parasites, guanylyl cyclases (GCs), which synthesise cyclic GMP (cGMP), are associated with a P4-ATPase-like domain in a unique bifunctional configuration. P4-ATPases generate membrane bilayer lipid asymmetry by translocating phospholipids from the outer to the inner leaflet. Here we investigate the role of *Plasmodium falciparum* guanylyl cyclase alpha (GCα) and its associated P4-ATPase module, showing that asexual blood stage parasites lacking both the cyclase and P4-ATPase domains are unable to egress from host erythrocytes. GCα-null parasites cannot synthesise cGMP, or mobilise calcium, a cGMP-dependent protein kinase (PKG)-driven requirement for egress. Using chemical complementation with a cGMP analogue and point mutagenesis of a crucial conserved residue within the P4-ATPase domain, we show that ATPase activity is up stream of and linked to cGMP synthesis. Collectively, our results demonstrate that GCα is a critical regulator of PKG and that its associated P4-ATPase domain plays a primary role in generating cGMP for merozoite egress.

## Introduction

Signalling through 3′, 5′-cyclic guanosine monophosphate (cyclic GMP or cGMP) regulates innumerable cellular processes across the animal kingdom, ranging from the control of cardiovascular function (reviewed in (Park et al., 2018)) and phototransduction (reviewed in (Michalakis et al., 2018)) in mammals through to differentiation and locomotion in several unicellular organisms (reviewed in (Baker and Kelly, 2004)). Two central players in the cGMP pathway are guanylyl cyclases (GCs), which synthesise cGMP from GTP, and phosphodiesterases (PDEs), which degrade cGMP. Cellular levels of cGMP are tightly regulated by the opposing action of these two enzyme classes. Upon reaching threshold levels, cGMP activates downstream effectors such as the cGMP-dependent protein kinase (PKG) or cGMP-gated ion channels. In the protozoan parasites responsible for malaria (*Plasmodium* spp.), PKG is thought to be the only direct effector of cGMP signalling since canonical cGMP-gated ion channels are absent from the genome (Gardner et al., 2002). Malaria parasites have two GC paralogues which are structurally distinct from both types of mammalian GC (Baker et al., 2017, Carucci et al., 2000, Linder et al., 1999). Each *Plasmodium* GC possesses a C-terminal domain with an overall topology similar to G protein-dependent adenylyl cyclases, comprising two catalytic domains each preceded by a set of six transmembrane helices. These twin catalytic domains contain all of the conserved amino acid residues required for cGMP synthesis. In addition, *Plasmodium* GCs also possess an N-terminal domain that has high structural similarity to P4-ATPases or flippases (Baker et al., 2017, Gao et al., 2018). In other organisms, flippases translocate phospholipids from the outer/luminal to the cytosolic leaflet of a membrane lipid bilayer, creating membrane asymmetry important for membrane remodelling, vesicular transport and signalling (reviewed in (Best et al., 2019, Lopez-Marques et al., 2011, Poulsen et al., 2008)). Most phospholipid flippases function in partnership with an integral membrane Cell Division Control protein 50 (CDC50) which acts as a chaperone for the enzyme and is required for flippase activity (Hiraizumi et al., 2019, Timcenko et al., 2019). The physical linkage of a guanylyl cyclase with a P4-ATPase domain is unique to *Plasmodium* and its apicomplexan and ciliate relatives (Baker et al., 2017, Bisio et al., 2019, Brown and Sibley, 2018, Gao et al., 2018, Gunay-Esiyok et al., 2019, Yang et al., 2019), but the functional significance of this unusual coupled architecture is unknown.

The complex life cycle of the malaria parasite is divided between a vertebrate host and a mosquito vector, and there is abundant evidence that cGMP signalling is crucial for parasite survival across multiple developmental stages. Whilst the asexual blood stages are solely responsible for clinical disease in the vertebrate host, a crucial step for transmission to the vector is the generation of sexual gametocyte forms. Upon uptake in a blood meal, the gametocytes are activated by a combination of reduced temperature with either an increase in pH or the presence of the mosquito factor xanthurenic acid (XA), (Billker et al., 1998) triggering the emergence of male and female gametes (gametogenesis). An early study using pharmacological agents suggested a role for cGMP signalling in male gamete activation in the mosquito (Kawamoto et al., 1990), and the action of XA was subsequently linked to elevated cGMP levels *in vitro* (Muhia et al., 2001). Using chemical genetic approaches, we showed that cGMP signalling through PKG is essential for gametogenesis (McRobert et al., 2008) and also for gliding motility of ookinetes (Brochet et al., 2014), motile forms that develop from the zygote that results from gamete fertilization. Of the two *P. falciparum* GC isoforms, GCβ is dispensable for blood stage development and is expressed exclusively in the mosquito stages (Taylor et al., 2008) where it is required for cGMP synthesis to drive and maintain *P. berghei* ookinete gliding motility (Brochet et al., 2014, Hirai et al., 2006, Moon et al., 2009). GCβ becomes polarised during ookinete formation, is stabilised by a CDC50 protein and is thought to elevate local levels of cGMP to facilitate gliding motility (Gao et al., 2018). The ookinetes penetrate the mosquito midgut epithelium and form oocysts in which thousands of sporozoites form; these then migrate to and invade the mosquito salivary glands. Dysregulation of cGMP levels by deletion of phosphodiesterase-gamma (PDEγ) in the rodent malaria species *Plasmodium yoelii* blocks salivary gland invasion and subsequent parasite development (Lakshmanan et al., 2015). In another rodent malaria parasite, *P. berghei*, PKG is required for sporozoite motility and invasion of hepatocytes following injection of sporozoites into a host by the mosquito (Govindasamy et al., 2016). Furthermore, conditional disruption of PKG reduces the release of merosomes containing merozoites into the bloodstream prior to erythrocyte invasion (Falae et al., 2010).

In contrast to GCβ, transcriptomic data indicate that *P. falciparum* GCα is expressed in both gametocytes and the asexual blood stages of the life cycle (https://plasmodb.org/plasmo/). The *GCα* gene has so far proved refractory to disruption in asexual blood stages (Kenthirapalan et al., 2016, Moon et al., 2009, Taylor et al., 2008), consistent with an important role during this clinically relevant stage. Confirmation of a key role for cGMP signalling in blood stages was obtained using a chemical genetic approach which demonstrated that PKG is essential for schizont rupture (Taylor et al., 2010). This observation was subsequently extended to show that just prior to merozoite egress, PKG regulates the discharge of the subtilisin-like protease SUB1 from organelles termed exonemes into the parasitophorous vacuole (PV) of mature *P. falciparum* schizonts, where it proteolytically processes a number of proteins required for merozoite egress and invasion (Collins et al., 2013b, Thomas et al., 2018). PKG activity is also required for the release of calcium ions (Ca^2+^) from internal stores that is a prerequisite for egress (Brochet et al., 2014). In efforts to dissect the mechanistic basis of this PKG-dependent egress pathway, comparative phosphoproteomic analysis identified 69 proteins that are phosphorylated in a cGMP-dependent manner (Alam et al., 2015). Very recently, the essential role for PKG in egress was confirmed through conditional genetic approaches (Koussis et al., 2020). Despite this clear evidence for a crucial role for PKG in the asexual blood stage life cycle, the role of GCα and its appended P4-ATPase domain remains unexplored.

Here we show that GCα is essential for *P. falciparum* blood stage egress and that GCα-null parasites cannot synthesise cGMP or mobilise Ca^2+^. Crucially, we show that activity of the P4-ATPase domain of GCα is essential and that it functions upstream of cGMP synthesis.

## Results

### *P. falciparum* GCα is expressed during late asexual blood stage development and localises to cytoplasmic vesicular structures in newly formed merozoites

Upon invasion of a red blood cell by a malaria merozoite, the parasite transforms through ring and trophozoite stages, and then undergoes DNA replication to form a multinucleated schizont that eventually segments to form a new generation of daughter merozoites. These are then released upon egress to invade fresh red cells and repeat the cycle. Invasion is driven by an actinomyosin-based contractile complex often referred to as the glideosome, which lies beneath the pellicular membrane of the merozoite. Data from several previous transcriptome studies show that GCα mRNA expression peaks at schizont stage, whilst proteomic analysis indicates expression in schizonts and merozoites (http://plasmodb.org/). To define the timing of expression of GCα at the protein level and to determine its subcellular localisation, we generated a *P. falciparum* line expressing GCα fused to a C-terminal triple hemagglutinin (3xHA) epitope tag. For this we used the *P. falciparum* 1G5 clone, which constitutively expresses a dimerisable Cre recombinase (DiCre) that can be activated by treatment with rapamycin (RAP) (Collins et al., 2013a). The tagging strategy used single crossover homologous recombination at the 3’ end of the endogenous *GCα* gene along with introduction of a human dihydrofolate reductase (hDHFR) selection cassette flanked by two *loxP* sites (Figure 1A). Initial selection for transformants with the antifolate WR99210, followed by several rounds of drug cycling, enriched for parasites in which integration had occurred. The *hDHFR* selectable marker was then recycled by RAP-induced Cre recombinase-mediated excision of the floxed sequence, leaving behind a single *loxP* site immediately downstream of the 3xHA epitope tag and translational stop codon (Figure 1A). Two parasite clones were obtained that showed sensitivity to WR99210, suggesting the expected genomic rearrangement with loss of the *hDHFR* selection marker (Figure 1B). The absence of the *hDHFR* cassette was confirmed in GCα:HA clone 1 by PCR analysis (Figure 1C), and this transgenic parasite clone (referred to as GCα:HA) was subsequently used in all further experiments.

**Figure 1.**
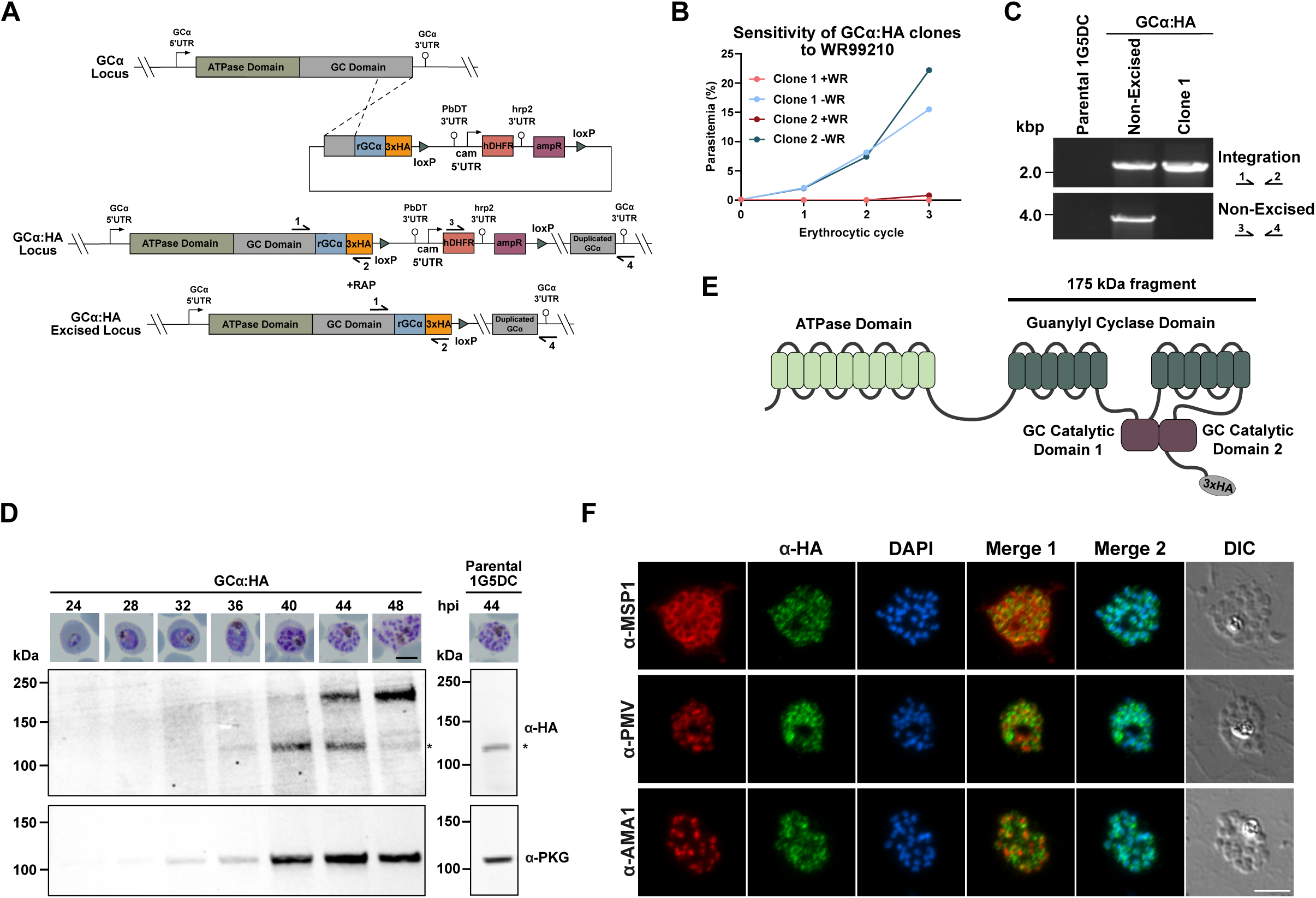
Generation of a GCα:HA-tagged line and spatio-temporal expression of GCα in *P. falciparum* blood stages. **(A)** Schematic representation of the single homology crossover approach used to fuse the 3’end of the endogenous *GCα* gene to a *3xHA* tag, and subsequent RAP-mediated excision of the *hDHFR* cassette. Promoters/5’ untranslated regions (UTRs) are indicated by arrows and 3’UTRs/terminators by lollipops. Triangles represent *loxP* sites, arrows with numbers represent the relative position of oligonucleotide primers used for diagnostic PCR. rGCα refers to recodonised *GCα* gene sequences. **(B)** Growth curve for two cloned GCα:HA lines sensitive to treatment with 2.5 nM WR99210, consistent with successful excision of the *hDHFR* cassette. Data presented are from counting parasites on Giemsa-stained blood smears. At least 100 parasites were counted per condition. Clone 1 was used for all further experiments. **(C)** Diagnostic PCR analysis confirming successful integration of the 3xHA tag, and efficient excision of the floxed *hDHFR* cassette in GCα:HA clone 1. Lane 1 Parental 1G5DC; lane 2 GCα:HA non-excised; lane 3 GCα:HA clone 1 (excised). **(D)** Western blot showing a time course of GCα:HA expression in *P. falciparum* blood stages. Parasites were harvested from cultures synchronised to a 2 h invasion window at the times indicated, with representative microscopy images shown above each sample (hpi, hours post invasion). Scale bar, 5 μm. Blots were probed with a monoclonal α-HA antibody to visualise the GCα:HA fusion protein and an anti-PKG antibody as a staging control. GCα:HA migrated as a ∼175 kDa fragment, while full length protein (predicted ∼499 kDa) could not be detected. Note that the additional band at ∼125 kDa (*) arises from a cross-reactivity of the α-HA antibody with an unrelated parasite protein, as it is also detected in extracts from the unmodified 1G5 parental *P. falciparum* line (right panel). **(E)** Schematic representation of the domain architecture of GCα, showing the N-terminal ATPase domain and the C-terminal guanylyl cyclase domain. The horizontal line with corresponding molecular mass shows the features likely contained in the C-terminal GCα-HA fragment detected by western blot. **(F)** Dual staining IFA analysis of mature GCα:HA schizonts. Formaldehyde-fixed thin films were stained with α-HA (green) and co-stained with antibodies to markers for known subcellular compartments (red): MSP1 (parasite plasma membrane, top panel), AMA1 (micronemes, middle panel), and plasmepsin V (endoplasmic reticulum, bottom panel). Scale bar, 5 μm. For additional images see Supplementary Figure 2A.

Tightly synchronised cultures of the GCα:HA line were sampled at 4 h intervals from early trophozoite stage (24 h post invasion) to mature schizont stage (48 h post invasion) and analysed by western blot using α-HA antibodies. This revealed a single specific signal migrating at ∼175 kDa that was most intense in mature schizonts (Figure 1D). This molecular mass is consistent with a C-terminal GCα-3xHA fragment predicted to comprise both the guanylyl cyclase catalytic domains (C1 and C2) plus the 12 associated transmembrane helices (Figure 1E). No signal was detectable at the ∼500 kDa mass expected for the full-length protein.

To confirm the timing of GCα expression at the single cell level and to reveal its subcellular localisation, we performed dual staining immunofluorescence microscopy (IFA) with anti-HA in combination with known markers for different subcellular compartments. In mature GCα:HA schizonts GCα localised to intracellular foci, but not the plasma membrane as established by co-staining with a merozoite surface protein 1 (MSP1) antibody (Figure 1F top panel). To further characterise the nature of the intracellular compartment occupied by GCα, we co-stained with antibodies that react with apical membrane antigen 1 (AMA1), a micronemal marker, or plasmepsin V, an ER resident protein. The α-HA staining showed no significant overlap with either of these markers, nor with a nuclear stain (Figure 1F middle and bottom panel). We conclude that GCα localises to non-apical, cytoplasmic vesicular structures and is maximally expressed in mature schizonts.

### GCα is essential for asexual blood stage growth and merozoite egress

To investigate the function and essentiality of GCα, we used the DiCre recombinase system (Collins et al., 2013a) to inducibly disrupt the *GCα* gene. For this, we created a GCα conditional knockout (cKO) line by introducing a second *loxP* site into the *GCα* locus of the GCα:HA line. This additional *loxP* site was incorporated within an artificial SERA2 intron (*loxPint*) (Jones et al., 2016) inserted into the ATPase domain of GCα using marker-free CRISPR/Cas9-mediated gene editing (Supplementary Figure 1A). Integration of the *loxPint* into the GCα coding region was confirmed by PCR in the uncloned parasite population (Supplementary Figure 1B) and following limiting dilution cloning two clones were obtained with the desired modification (Supplementary Figure 1C). One of these clones, called GCα:HA:cKO, was used in all further experiments.

Activation of Cre recombinase in the GCα:HA:cKO line by addition of RAP was expected to excise DNA sequences encoding part of the ATPase domain and the entire cyclase domain of GCα, abrogating both enzymatic activities (Figure 2A). To examine the efficiency of excision, synchronous cultures of young ring stage parasites were treated with either 50 nM RAP or the equivalent concentration of vehicle only (DMSO) for 2 h and allowed to mature to schizont stage. PCR analysis showed highly efficient RAP-induced excision (Figure 2B) whilst western blot analysis showed complete loss of the 175 kDa GCα-3xHA signal in the RAP-treated GCα:HA:cKO schizonts, confirming efficient gene disruption (Figure 2C). In addition, immunofluorescence microscopy showed loss of GCα-3xHA at the single cell level (Figure 2D and Supplementary Figure 2B and 2C).

**Figure 2.**
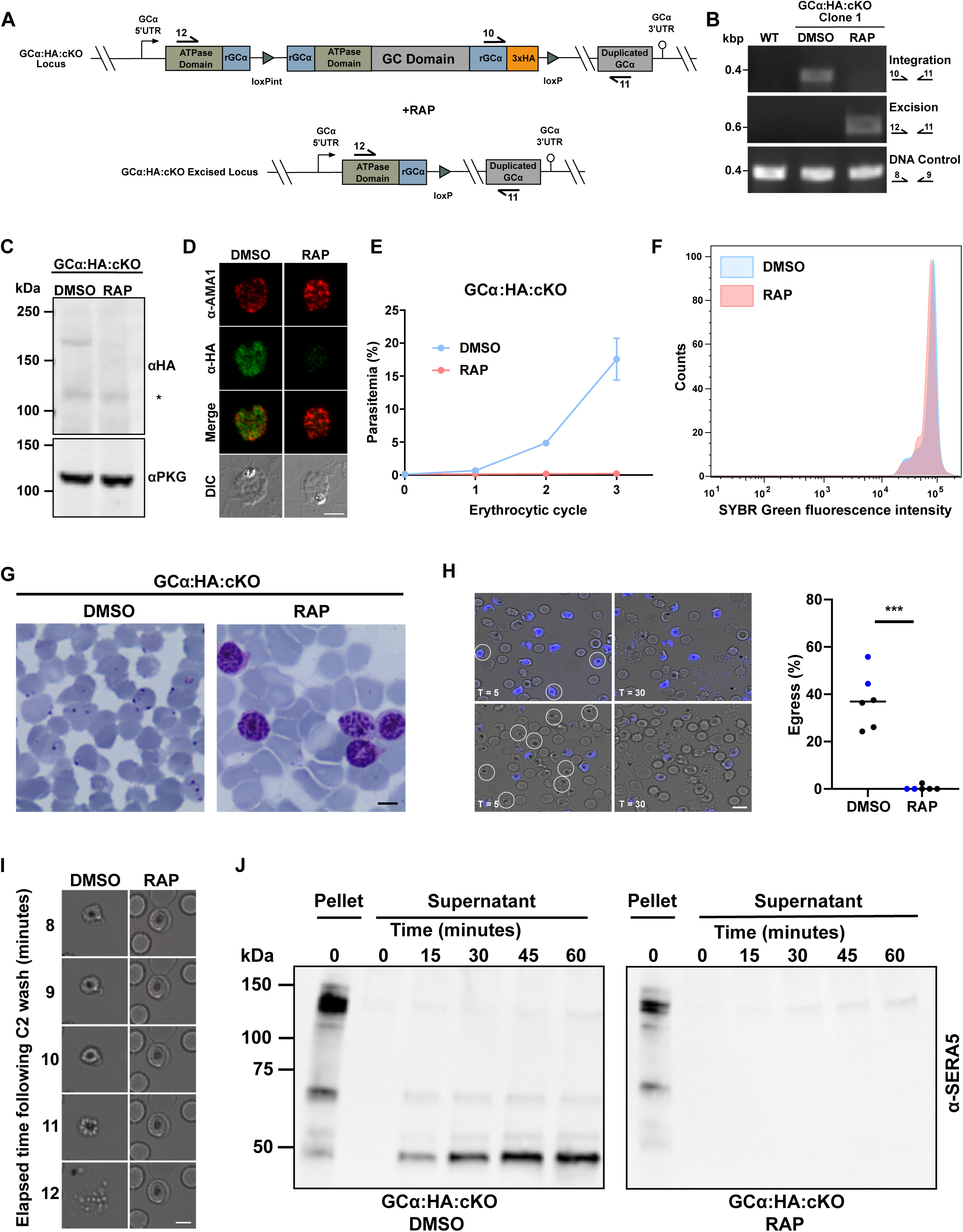
Efficient conditional disruption of GCα results in a complete egress block. **(A)** Schematic representation of RAP-induced excision of a portion of the ATPase domain and the entire GC domain of GCα in GCα:HA:cKO. Arrows represent the relative position of oligonucleotide primers used for diagnostic PCR screens. **(B)** Diagnostic PCR analysis of DMSO- and RAP-treated GCα:HA:cKO parasites showing efficient excision of the *GCα* gene following RAP treatment. A DNA control PCR was included, which amplified a small segment at an independent locus (Pf3D7_1342600) to confirm the quality of the DNA used. Specific PCR primers used are indicated on the right and their binding sites are shown in (A). **(C)** Western blot analysis of DMSO- and RAP-treated GCα:HA:cKO schizont lysates probed with an α-HA antibody, showing loss of the ∼175 kDa band in the RAP-treated sample, while the non-specific ∼125 kDa (*) band can still be observed in both samples. The blots were also probed with an α-PKG antibody to serve as a loading control. **(D)** Parallel IFA analysis of mature schizonts from DMSO- and RAP-treated GCα:HA:cKO cultures shows loss of GCα-3xHA protein at the single cell level. Formaldehyde-fixed thin films were probed with α-HA (green) and α-AMA1 (red) antibodies. Scale bar, 5 μm. This analysis also confirms the specificity of the signal obtained with anti-HA. Additional images are shown in Supplementary Figure 2B and 2C. **(E)** Growth curves showing parasitaemia of GCα:HA:cKO cultures treated with either DMSO or RAP, measured over three replication cycles by flow cytometry-based counting of SYBR Green positive cells. Data points plotted are means from two repeat experiments, each performed in triplicate. Error bars, S.D. Note that from ∼16 days post RAP treatment parasites emerged, but these were evidently not GCα-null since PCR showed that the GCα locus was intact (Supplementary Figure 3A and 3B). **(F)** Comparison of the DNA content in wild type and GCα-null schizonts. Schizonts obtained from synchronous DMSO- and RAP-treated GCα:HA:cKO cultures were arrested using compound 2 to prevent egress, fixed at ∼46 h post invasion and parasite DNA stained with SYBR Green. SYBR Green fluorescence intensity was measured by FACS counting from technical triplicates. **(G)** Representative microscopy images of Giemsa-stained parasites from DMSO- and RAP-treated GCα:HA:cKO cultures at ∼50 h post invasion, showing the accumulation of schizonts in the RAP-treated culture while new ring stage parasites formed in the DMSO control culture. Scale bar, 5 μm. **(H)** Combined DIC and fluorescence images from time-lapse video microscopy of DMSO- and RAP-treated GCα:HA:cKO schizonts taken at 5 minutes (T=5) and 30 minutes (T=30) after release from a compound 2 block applied to transiently prevent and subsequently synchronise egress. In each experiment, one subset of parasites was pre-treated with Hoechst to stain the nuclei so that DMSO- and RAP-treated parasites could be viewed simultaneously in the same imaging chamber. In the top panel, DMSO-treated parasites are Hoechst-stained, while in the bottom panel, RAP-treated parasites are Hoechst-stained. Schizonts visible the first frame that rupture over the course of the video are circled in white. Scale bar, 10 μm. The graph on the right shows a quantification of the percentage of DMSO- and RAP-treated schizonts that egressed in each 30 minute video. Data were collected from 6 videos, with Hoechst-treated samples depicted in blue. Statistical significance was measured by unpaired t-test, where *** signifies p < 0.0001. **(I)** Time series of individual stills from GCα:HA:cKO time-lapse video microscopy in (G). Images following representative schizonts for each condition (DMSO and RAP) from 8 to 12 minutes post compound 2 washout are shown. Note that the PVM around the RAP-treated schizont remains intact throughout the time series, while the DMSO control has completed egress by 12 min. Scale bar, 5 μm. **(J)** Western blot analysis monitoring the release of SERA5 into the culture supernatant of DMSO- and RAP-treated GCα:HA:cKO schizonts, as a measurement of egress over time. Sampling times are indicated in minutes. Pellet samples were included as loading controls.

To assess the impact of GCα disruption on parasite viability and growth, we used flow cytometry to assess replication assay of DMSO-treated (control) and RAP-treated GCα:HA:cKO parasites over 3 erythrocytic growth cycles. This revealed a complete growth arrest resulting from disruption of the GCα locus (Figure 2E). Examination of Giemsa-stained GCα:HA:cKO parasites in the cycle of gene excision (cycle 0) revealed that RAP-treated cultures developed normally and were able to form mature segmented schizonts with a DNA content indistinguishable from control schizonts (Figure 2F). However, upon further incubation we observed an accumulation of schizonts and a complete absence of newly formed ring stage parasites, indicating that GCα is required for egress (Figure 2G). To further characterise and quantify the egress defect in the GCα-null parasites, highly synchronised mature cycle 0 schizonts from DMSO- and RAP-treated cultures were monitored by live time-lapse microscopy. This confirmed that GCα-null parasites were unable to egress (Figure 2H and Supplementary Videos). Close examination of individual frames of the time-lapse video series revealed no signs of swelling or PVM rupture in GCα-null schizonts (Figure 2I). Consistent with the above results, the abundant PV-resident protein SERA5, which is released during egress, was not detectable in culture supernatant samples from RAP-treated cultures (Figure 2J). Collectively, these results establish that GCα plays an essential role in the asexual blood stage life cycle and is required for merozoite egress.

### GCα is responsible for cGMP production and essential for calcium release in asexual blood stage schizonts

The phenotypic similarity between the egress block observed in GCα-null schizonts and that produced by the PKG inhibitor compound 2 (Figure 2I and Figure 3A) is consistent with GCα-null schizonts failing to activate PKG due to a lack of cGMP synthesis. To evaluate the impact of GCα disruption on cyclic nucleotide production, we compared intracellular cGMP concentrations in mature cycle 0 GCα:HA:cKO schizonts from DMSO- or RAP-treated cultures. For these experiments, in order to prevent egress of the DMSO-treated (control) schizonts, both sets of parasites under comparison were maintained in the presence of compound 2. This analysis showed that ablation of GCα led to a 94.5% reduction in cGMP levels compared to control parasites (Figure 3B). Treatment with the PDE inhibitor zaprinast, which blocks cGMP hydrolysis in *P. falciparum* schizonts (Collins et al., 2013b) and thus artificially elevates cGMP levels, resulted in a significant increase in cGMP levels in the DMSO- but not RAP-treated samples, confirming that GCα-null schizonts are unable to synthesise cGMP (Figure 3B). Importantly, we detected no significant differences in cAMP levels between the control and GCα-null parasites, nor did cAMP levels increase upon zaprinast treatment (Figure 3C) consistent with our previous findings (Collins et al., 2013b).

**Figure 3.**
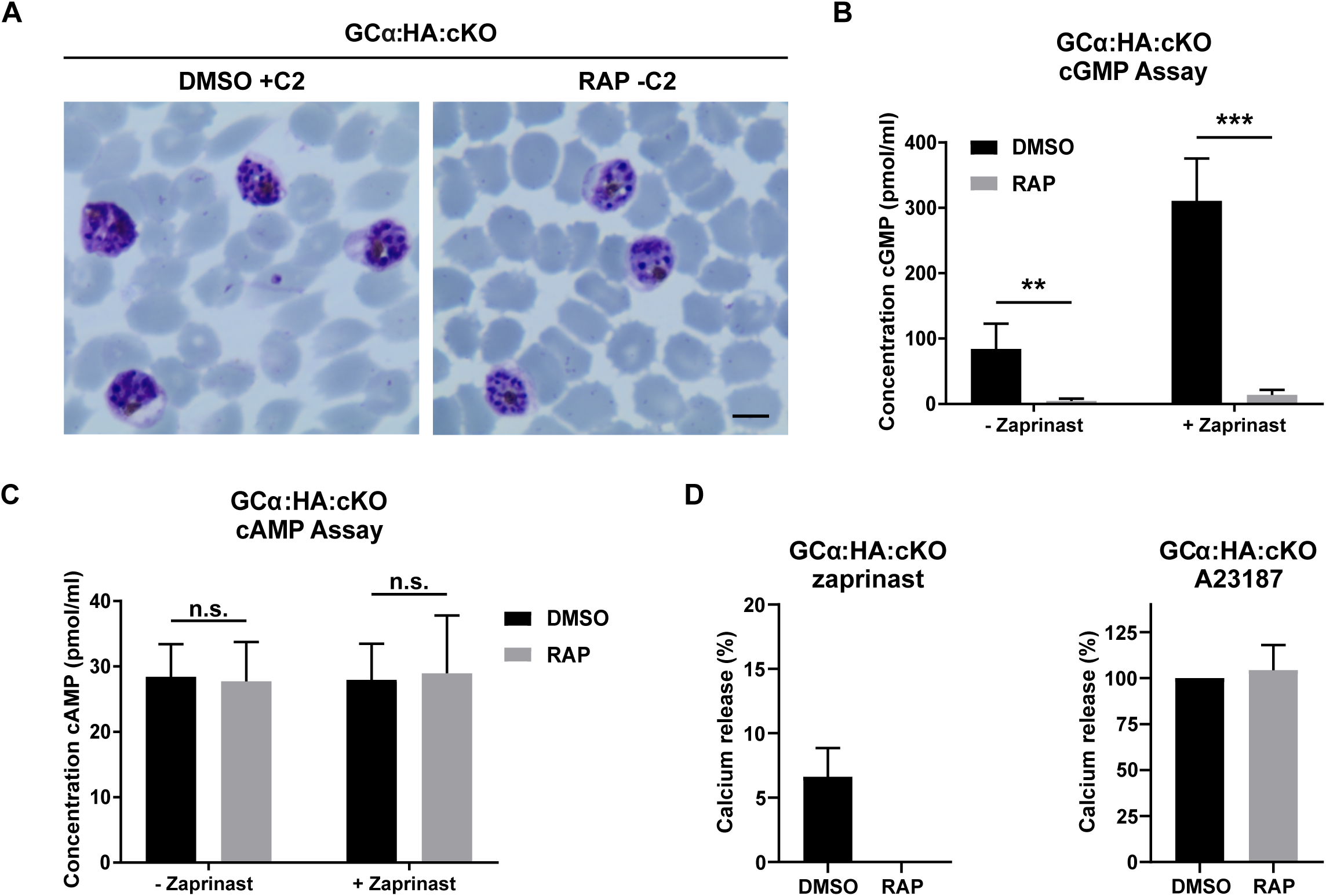
GCα disruption leads to a complete loss of cGMP production and a block in PKG-mediated calcium release. **(A)** Representative images of Giemsa-stained schizonts from DMSO-treated GCα:Ha:cKO parasites in the presence of 1.5 μM compound 2, and RAP-treated GCα:HA:cKO parasites at 50 h post invasion. PKG inhibition leads to a similar phenotype to that observed in GCα-null parasites. Scale bar, 5 μm. **(B)** Comparison of intracellular cGMP levels in wild type and GCα-null schizonts. Schizonts from DMSO- and RAP-treated GCα:HA:cKO cultures were matured in presence of 1.5 µM compound 2 to prevent egress. Segmented schizonts were either lysed directly (- zaprinast) or pre-treated with 75 μM zaprinast for 3 minutes prior to cell lysis (+ zaprinast). All samples were lysed in 0.1 M HCl to inactivate all PDEs and extracts analysed using a commercial ELISA-based cGMP detection assay. Data presented are means from four independent experiments, each assayed in duplicate. Error bars, S.D. Statistical significance was measured by unpaired t-test, ** signifies p < 0.01 whereas *** signifies p < 0.001. **(C)** The samples described in (B) were assayed in parallel for cAMP with an ELISA-based cAMP kit from the same manufacturer. Data presented are means from four independent experiments, each performed in duplicate. Error bars, S.D. Statistical significance was measured by unpaired t-test, n.s. indicates not significant. **(D)** Determination of PKG-mediated calcium release from internal stores in wild type and GCα KO schizonts. Fluo-4-loaded mature DMSO- and RAP-treated GCα:HA:cKO schizonts were treated with 75 μM zaprinast (left panel) or 20 μM ionophore A23187 (right panel) and the mobilisation of Ca^2+^ measured by fluorimetry. Zaprinast signals (left plot) were normalised to their respective ionophore control. Ionophore signals were normalised to the DMSO-treated sample (right plot). Data plotted are mean values from two independent experiments performed in triplicate. Error bars, S.D.

Next, we investigated whether Ca^2+^ release from internal stores into the cytosol, a PKG-dependent process essential for merozoite egress (Brochet et al., 2014), was impaired in the GCα knockout parasites. Treatment with the PDE inhibitor zaprinast specifically induces PKG-mediated calcium release (Brochet et al., 2014). We measured zaprinast-induced Ca^2+^ release in DMSO- and RAP-treated GCα:HA:cKO parasites by loading cells with the fluorescent calcium indicator Fluo-4 AM. As shown in Figure 3D, while zaprinast treatment stimulated elevated cytosolic Ca^2+^ levels in control schizonts no such elevation of Ca^2+^ levels was detected in the GCα-null schizonts (Figure 3D). Importantly, both control and GCα-null schizonts showed similar response levels to the calcium ionophore A23187, which allows Ca^2+^ ions to cross cell membranes, indicating that internal calcium stores were not affected in the absence of GCα (Figure 3D). Together, these results clearly identify GCα as a functional GC and establish that it generates the cGMP signal in asexual blood stage schizonts essential for PKG activation, PKG-dependent Ca^2+^ release and ultimately merozoite egress.

### Chemical complementation of GCα-null parasites with PET-cGMP rescues the egress defect

Since the egress phenotype observed in GCα-deficient parasites is most likely due to their inability to produce cGMP to activate PKG, we reasoned that bypassing the need for cGMP synthesis by directly activating PKG could potentially rescue the egress defect. We also anticipated that this might reveal a second phenotype resulting from loss of the ATPase domain. To test this notion, we supplemented cultures containing highly mature RAP-treated cycle 0 GCα:HA:cKO schizonts with either cGMP or two cGMP analogues, 1-NH_2_-cGMP and PET-cGMP, both of which have previously been shown to activate recombinant apicomplexan PKG (Byun et al., 2020, Salowe et al., 2002). While cGMP and 1-NH_2_-cGMP show similar levels of membrane permeability, PET-cGMP is 50-fold more lipophilic (https://www.biolog.de/technical_info_lipophilicity-data) due to its possession of an etheno phenyl extension to the purine ring of cGMP. Following a 1 h incubation in the presence of various concentrations of cGMP, 1-NH_2_-cGMP or PET-cGMP, microscopic examination of the cultures revealed that PET-cGMP was highly effective at rescuing the egress block, resulting in the release of merozoites at all concentrations tested (Figure 4A and Supplementary Figure 4A). While rings with normal morphology could be observed in the cultures containing the lower concentrations of PET-cGMP (15, 31.2 and 62.5 μM), the cultures supplemented with higher PET-cGMP concentrations contained numerous intracellular or extracellular pyknotic forms, indicating toxicity (Figure 4B). Few rings were observed in the cGMP and 1-NH_2_-cGMP-treated cultures, indicating limited activity of these compounds (Figure 4A). It was concluded that PET-cGMP efficiently reversed the egress defect displayed by the GCα-deficient parasites (Figure 4A).

**Figure 4.**
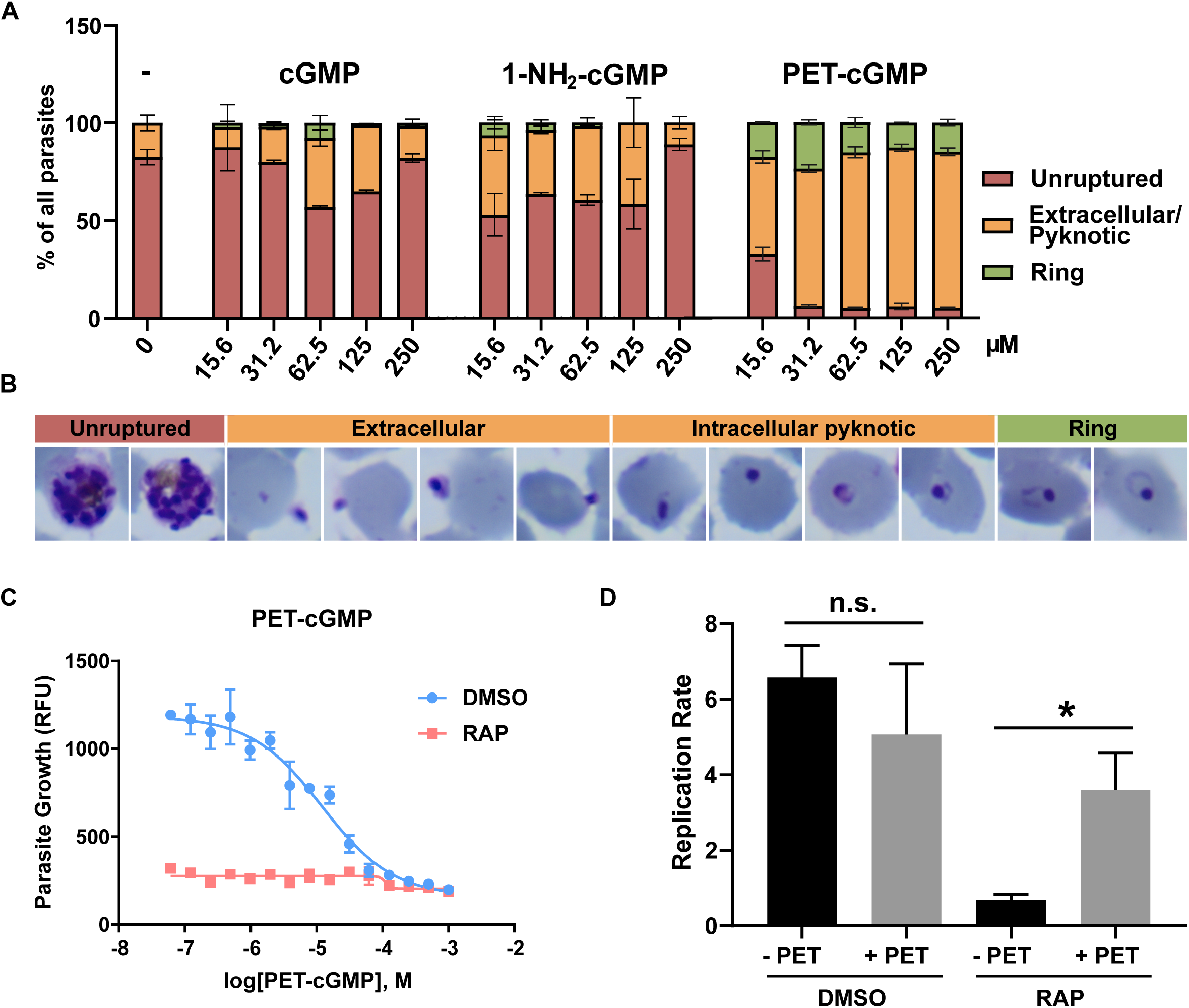
Addition of PET-cGMP efficiently rescues the egress block phenotype of GCα knockout schizonts. **(A)** Quantification of the effect of various concentrations of cGMP, 1-NH_2_-cGMP and PET-cGMP on the reversal of the egress block and promotion of ring stage development in RAP-treated GCα:HA:cKO parasites. Ten microscopic fields per condition were counted blind by two researchers and parasites scored as either unruptured (trophozoites and schizonts), uninvaded (free merozoites and pyknotic), or invaded (ring stages). Data are presented as proportions of the different forms of total parasites counted. Values are means of counts by each researcher. Error bars, S.D. **(B)** Representative Giemsa images of pyknotic forms observed in smears taken from RAP-treated GCα:HA:cKO cultures 1 h following addition of 62.5 μM PET-cGMP in the experiment quantified in (A). **(C)** Continuous PET-cGMP treatment is toxic and does not rescue GCα-null parasite growth. Growth of DMSO- and RAP-treated GCα:HA:cKO parasites at different concentrations of PET-cGMP as measured in a 72 h SYBR Green growth assay. Assays were repeated twice and the mean EC_50_ of PET-cGMP was measured as 12.49± 0.06 μM in wild type parasites (DMSO). Error bars. S.D. **(D)** Comparison of replication rates from cycle 0 to cycle 1 of DMSO- and RAP-treated GCα:HA:cKO parasites cultured in the presence or absence of 30 μM PET-cGMP, as measured by flow cytometry-based counting of SYBR Green-positive cells. PET-cGMP was added when mature segmented schizonts appeared in culture and washed off ∼10 h later, once rings had formed. Data points plotted are means from two repeat experiments, each performed in triplicate. Error bars, S.D. Statistical significance was measured by a ratio paired t-test with * signifying p > 0.05 (0.0177). n.s., not significant.

To determine the optimum concentrations of PET-cGMP to sustain replication of the GCα-null parasites, parasite proliferation was assessed over a 72 h period (1.5 erythrocytic cycles) in the continuous presence of various concentrations of PET-cGMP. Interestingly, this revealed that none of the concentrations tested could rescue replication of the GCα-null parasites (Figure 4C), including those concentrations that efficiently rescued egress in the short-term assays described above. To try to understand this, we performed parallel assays assessing growth of control, DMSO-treated GCα:HA:cKO parasites in cultures supplemented with the same range of PET-cGMP concentrations. This revealed that PET-cGMP is toxic, with an EC_50_ of 12.49±0.06 μM (Figure 4C). Given its capacity to rescue the egress defect in GCα-null parasites, we considered that the toxic effect of prolonged exposure to the compound is likely explained by premature activation of PKG as the parasites matured, perhaps resulting in premature egress and release of non-invasive merozoites, a phenomenon previously observed upon treatment of schizonts with zaprinast (Collins et al., 2013b). In support of this model, further work showed that mature DMSO- and RAP-treated GCα:HA:cKO schizonts tolerated short-term incubation with 30 μM PET-cGMP followed by wash off of the compound ∼10 h later, after the majority of schizonts had ruptured and formed rings; under these conditions, the new rings successfully matured to form cycle 1 schizonts (Supplementary Figure 4B). Based on these findings, we used flow cytometry to quantify the degree of rescue that could be achieved over a single egress/invasion cycle. Quantification of cycle 1 parasite levels the following day showed that the presence of PET-cGMP during egress and invasion produced a 3.6-fold increase in parasitaemia in the GCα-null cultures, whilst parallel cultures of GCα-null parasites lacking PET-cGMP completely failed to expand (Figure 4D). In contrast, the replication rate of control parasites was slightly reduced (from 6.5 to 5-fold) by similar short-term treatment with PET-cGMP, perhaps indicative of the low levels of toxicity of the compound (Figure 4D).

These results confirmed the capacity of PET-cGMP to rescue the egress defect in the GCα-null parasites. Of particular significance, since the conditional strategy used to disrupt the *GCα* gene was designed to excise key segments of both the cyclase and ATPase-encoding sequences, the results indicated that either the ATPase domain has no essential function, or that it acts upstream of cGMP synthesis allowing its role to be bypassed by the presence of the cGMP analogue.

### A conserved catalytic Asp residue within the ATPase domain is required for parasite survival

The N-terminal portion of GCα encodes a putative P-type ATPase that shares closest homology with type IV ATPases (P4-ATPases) (Baker et al., 2017) which in other organisms flip phospholipids from the outer to the inner leaflet of a lipid bilayer (Hiraizumi et al., 2019, Timcenko et al., 2019). P-type ATPases possess 10 transmembrane helices that facilitate transport of ligands according to the Post-Albers mechanism (Lutsenko and Kaplan, 1995). This requires the following cytoplasmic components: a nucleotide binding domain (N-domain), which binds ATP; a phosphorylation domain (P-domain), which contains a highly conserved aspartate (Asp) residue that becomes autophosphorylated to form an aspartyl phosphate intermediate; and an actuator domain (A-domain), which dephosphorylates the phosphorylation domain (Kuhlbrandt, 2004, Montigny et al., 2016) (Figure 5A). ATP-dependent autophosphorylation of the conserved Asp, which lies within a DKTGT motif, leads to a rotational change in the actuator domain surrounding the phosphorylation site; dephosphorylation is then coupled to inward transport of phospholipids or ions in the respective P-type ATPase families. Alignment of the GCα N-terminal region with other P4-ATPases, including human ATP8A1 for which a crystal structure has recently been determined (Hiraizumi et al., 2019) shows that of the 56 residues identical in P4-ATPases and P2-ATPases (a Ca^2+^ transporting SERCA and a Na^+^-K^+^ P-type ATPase) that are required for ATP binding and catalysis, 47 are also identical in GCα and 2 are similar, fully consistent with the GCα sequence representing a functional ATPase domain (Supplementary Figure 5).

**Figure 5.**
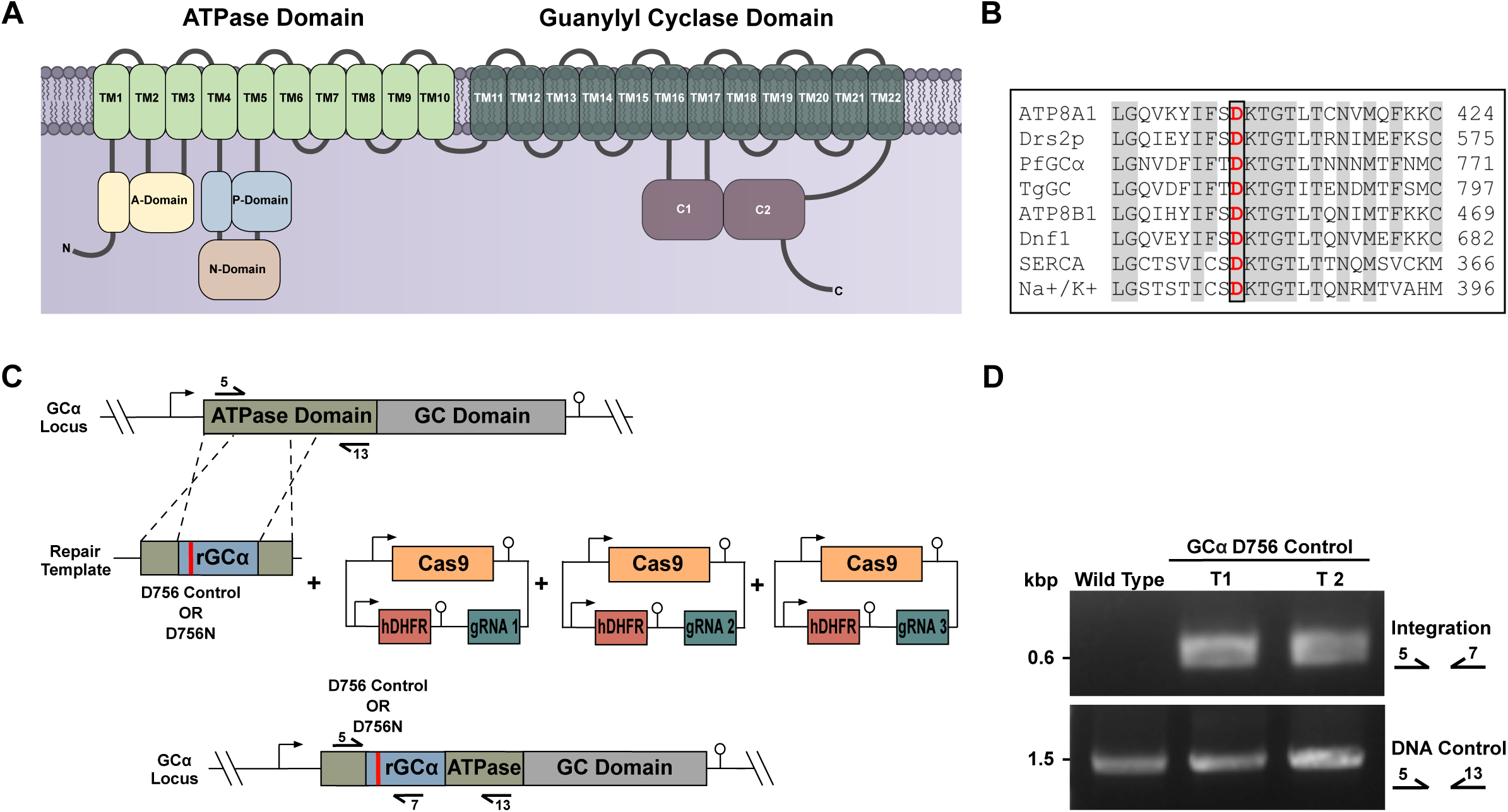
Key functional domains of the ATPase domain of GCα and strategy for substitution of the phosphorylation site aspartate. **(A)** Cartoon showing details of the predicted ATPase topology of *P. falciparum* GCα. **(B)** Amino acid alignment of the phosphorylation site within the P-domain of P4-ATPases. The aligned sequences are: GCα (PlasmoDB identifier: PF3D7_1138400), human ATP8A1, UniProt: Q9Y2Q0-2), human ATP8B1 (UniProt: O43520), yeast Drs2p (UniProt: P39524), yeast Dnf1 (UniProt: P32660), *T. gondii* GC (accession number EPT31724), *P. falciparum* GCβ (PlasmoDB identifier: PF3D7_1360500), rabbit SERCA Ca^2+^-ATPase isoform 1a (UniProt: P04191) and dogfish Na^+^/K^+^-ATPase (UniProt: Q4H132). **(C)** Schematic representation of the marker-free CRISPR/Cas9-mediated approach used to introduce synonymous or non-synonymous mutations into the ATPase domain of *GC*α in the GCα:HA line. Arrows with numbers represent the relative position of oligonucleotide primers used for diagnostic PCR. A pool of three Cas9 plasmids harbouring different gRNA cassettes was transfected along with a linearised plasmid containing synonymous mutations (D756 control) or mutations resulting in an amino acid change (D756N), flanked by recodonised (rGCα) and endogenous GCα sequences serving as template for homology-directed repair. Promoters/5’UTRs and 3’UTRs/terminators are indicated by arrows and lollipops, respectively. **(D)** Diagnostic PCR evidencing modification of the GCα locus by integration of the D756 control construct in two independent transfections. PCR primers used are indicated on the right and their respective binding sites shown in (C).

Previous functional investigations (reviewed in (Best et al., 2019) and two recent structural studies on yeast Drs2p and human ATPase8A1 in complex with their respective CDC50 partners (Hiraizumi et al., 2019, Timcenko et al., 2019), have demonstrated the importance of the transmembrane domains of P4-ATPases, particularly TM1-4 (but also TM6), for lipid binding and translocation. Key amongst these is the PISL motif in TM4 which is highly conserved in P4-ATPases and is also present in the GCα sequence. This motif constitutes an important difference between the phospholipid and cation transporting ATPases and is crucial for binding and translocation of their respective substrates. P4-ATPases have a P<u>IS</u>L motif whereas the ion transporters have a P<u>EG</u>L motif. GCα has a P<u>IS</u>I motif at this position which contains the crucial IS pair diagnostic of phospholipid binding/translocation rather than cation binding/translocation (Supplementary Figure 5). The QQ motif at the C-terminus of TM1 contains non-charged polar residues (e.g. Q and N) which in human ATP8A1 determine selectivity for PS in P4-ATPases. Q and N can hydrogen bond to the negative charge of the polar head group of PS, whereas the presence of small non-polar residues (A and G) in place of the QQ motif preclude hydrogen bonding to PS and are features of P4-ATPases that translocate PC (Hiraizumi et al., 2019). GCα has a QN pair corresponding to the QQ motif of human ATP8A1 which is compatible with flipping of PS.

To address whether the ATPase domain of GCα is required for blood stage replication, we investigated whether parasites could replicate following substitution of the conserved Asp (D756) in a manner designed to block formation of the aspartyl phosphate intermediate and thus ablate any enzymatic activity of the ATPase. To do this, we employed marker free CRISPR/Cas9-mediated gene editing to introduce an Asn substitution of GCα D756 (Figure 5B) (Figure 5C and Supplementary Figure 1). In parallel control manipulations, we used a similar approach to introduce synonymous mutations that did not alter the amino acid sequence but that could be differentiated from the wild type locus at the nucleotide level. While parasites with a modified locus appeared in two out of two independent transfections in these control cultures by two weeks post transfection (Figure 5D), no parasites emerged from three independent D756N transfections performed in parallel (one was performed in parallel with the successful integration of the *loxPint*) even following extended culture (6 weeks). Our failure to obtain viable parasites harbouring the D756N mutation after three attempts, whilst readily obtaining transgenic parasites possessing synonymous mutations of the D756 codon, strongly suggests that an active ATPase domain is required for parasite survival. These data, combined with our ability to rescue the GCα-null growth phenotype by chemical complementation with PET-cGMP, indicate that the ATPase domain acts upstream of and is required for GC activity and that the ultimate role of the ATPase domain is to facilitate cGMP synthesis.

## Discussion

Although the single GC in the related apicomplexan parasite *T. gondii* (Bisio et al., 2019, Brown and Sibley, 2018, Gunay-Esiyok et al., 2019, Yang et al., 2019) and GCβ in *P. falciparum* (Taylor et al., 2008), *P. berghei* (Hirai et al., 2006, Moon et al., 2009) and *P. yoelii* (Gao et al., 2018) have been previously analysed by reverse genetics, to date there have been no functional studies of GCα in the asexual blood stages of *Plasmodium*. Here we have made important advances in understanding the essentiality and function of this key component of the cGMP signalling pathway.

Epitope tagging of the endogenous *P. falciparum* GCα locus indicated that the protein is located in puncta within the confines of individual merozoites within mature schizonts. We propose that these puncta represent cytoplasmic membranous structures, consistent with the architecture of the protein which has 22 predicted transmembrane helices. It is likely that the GCα topology is orientated such that the twin GC catalytic domains face the cytosol in order to allow synthesis of cGMP to activate PKG, the majority of which is cytosolic (Hopp et al., 2012). Our observed localisation for *P. falciparum* GCα is reminiscent of that recently described for GCα in *P. yoelii* gametocytes (Jiang et al., 2020). In contrast, in *Toxoplasma* (Bisio et al., 2019, Brown and Sibley, 2018, Gunay-Esiyok et al., 2019, Yang et al., 2019) the single GC has been localised to the plasma membrane, primarily at the apical pole, in extracellular parasites.

Epitope-tagged *P. falciparum* GCα migrated in western blots as a ∼175 kDa species, much smaller than the predicted full-length protein (∼500 kDa). The signal therefore likely represents a C-terminally truncated proteolytic fragment resulting from protein instability during detergent extraction. A similar phenomenon has also been observed in *P. yoelii*, where GCα migrates at ∼175 kDa instead of the predicted full-length ∼450 kDa (Jiang et al., 2020), as well as in *T. gondii,* where TgGC predominantly migrates at ∼125 kDa and ∼75 kDa, with only small amounts of the ∼ 460 kDa species corresponding to full-length protein detected in some studies (Bisio et al., 2019, Brown and Sibley, 2018, Jia et al., 2017, Yang et al., 2019). Given the evidence suggesting that the *Toxoplasma* GC needs to be full length to be functional, it seems likely that full-length *P. falciparum* GCα protein is present but below detection limits in our experiments.

Conditional deletion of a large segment of GCα spanning both the GC catalytic domain and the ATPase-like domain prevented asexual blood stage replication, with a selective block in egress. This finding was in contrast to results generated in a previous global transposon-based gene knockout study which suggested that GCα was dispensable for blood stage replication (Zhang et al., 2018). However, that study acknowledged that false negatives can be obtained using their approach depending upon the position of transposon insertion. In an *in vivo* global gene knockout study in *P. berghei*, parasites in which GCα was disrupted grew slowly and were close to the cut off for those essential for blood stage replication (Bushell et al., 2017) which is more consistent with our findings in *P. falciparum*.

We have previously reported that GCα possesses all the conserved amino acid residues required for catalytic activity (Carucci et al., 2000). However, the ability of GCα to synthesise cGMP had previously not been demonstrated. The lack of detectable cGMP generation in GCα-null parasites confirms that it is a functional GC and also indicates that GCβ, which is expressed only during mosquito stage development, cannot compensate for the absence of GCα in blood stages. The lack of cGMP synthesis fully explains the egress phenotype of GCα-null parasites, because activation of PKG by cGMP is known to be essential for egress (Collins et al., 2013b, Taylor et al., 2010). PKG activity also induces calcium release from internal stores in *P. falciparum* schizonts which is required for egress. This is concordant with our finding that GCα-null schizonts did not mobilise calcium in response to cGMP elevation by the PDE inhibitor zaprinast. Together, these results establish that GCα is the key regulator upstream of PKG activity in *P. falciparum* blood stages. Disruption of the *Toxoplasma* GC also abolishes the rises in cytosolic Ca^2+^ required for secretion of micronemal proteins (Yang et al., 2019) and motility (Moudy et al., 2001, Roiko et al., 2014).

It has been suggested that the chemical gradient of phospholipids generated by P4-ATPases is akin to the chemical gradient of ions created by P2-ATPases to mediate signal transduction (Roland and Graham, 2016). To investigate whether the ATPase-like domain in PfGCα is likely to be a catalytically active ATPase we attempted to generate parasites possessing a substitution of a highly conserved aspartate (D756) that is autophosphorylated in functional ATPases. These parasites could not be selected whilst those engineered to reconstitute the wild type aspartate proliferated normally, providing evidence that the ATPase domain is catalytically active and serves an essential function. Our finding that the GCα-null phenotype could be rescued by addition of the membrane-permeable cGMP analogue PET-cGMP, strongly suggests that the function of the ATPase domain is upstream of and directly related to cGMP synthesis. Our findings are consistent with previous studies in *Toxoplasma* where the ATPase domain was also shown to be critical for GC function (Bisio et al., 2019, Brown and Sibley, 2018, Yang et al., 2019). Complementation of GC-null *Toxoplasma* with a panel of mutants demonstrated that the ATPase domain is required for trafficking of GC, activity and maximal GC function since *stimulated* micronemal secretion was only partially reduced in the ATPase D728A mutant whilst natural microneme secretion was prevented (Brown and Sibley, 2018).

Although our data suggest that the activity of the ATPase domain of GCα is an essential upstream regulatory factor for cGMP synthesis, our study does not further address its biochemical activity or indeed how it facilitates cGMP synthesis. The single *Toxoplasma* GC is involved in sensing phosphatidic acid as well as changes in pH and potassium levels to mediate egress (Bisio et al., 2019, Yang et al., 2019). A recent suggestion that *P. falciparum* GCα might flip phosphatidylcholine (Paul et al., 2020) to mediate cGMP-stimulated egress was not supported by direct evidence of GCα-mediated flipping of phosphatidylcholine. Our analysis has shown the presence of a QN pair in *P. falciparum* at the C-terminus of TM1 in GCα, which corresponds to the QQ motif of human ATP8A1 (Hiraizumi et al., 2019), suggesting that the sequence is compatible with flipping of PS. Future work will be needed to determine whether regulation of cGMP synthesis by GCα requires flipping or sensing of phospholipids by the ATPase domain and whether generation of lipid asymmetry across membranes contributes to activation of cGMP synthesis.

Synthesis of cGMP by GCα has been linked to the stimulation of gametogenesis by xanthurenic acid (XA) (Muhia et al., 2001), but it is not clear whether XA stimulates GCα activity directly or indirectly through other protein mediators. However, a study in *P. yoelii* has recently identified a protein called GEP1 that interacts with GCα, showing that both are required for XA-stimulated gametogenesis (Jiang et al., 2020). Although cGMP synthesis and PKG activation are required for merozoite egress, the nature of the upstream signal that activates GCα in *Plasmodium* blood stages is unknown. Just prior to natural egress in *Toxoplasma*, the parasitophorous vacuole is acidified which triggers micronemal secretion. A similar mechanism may operate in *Plasmodium*. Future work will be needed to determine the events that occur upstream of cGMP signalling in blood stages and to understand the role of the P4-type ATPase domain in mediating the egress signal.

## Materials and Methods

### Small molecules and antibodies

WR99210 was kindly provided by Jacobus Pharmaceuticals (New Jersey, United States). The PKG inhibitor compound 2 (4-[7-[(dimethylamino)methyl]-2-(4-fluorphenyl)imidazo[1,2-*α*]pyridine-3-yl]pyrimidin-2-amine) was synthesised by MRC Technology (London, United Kingdom). Rapamycin, calcium ionophore A23187, the cysteine protease inhibitor E64 and the PDE inhibitor zaprinast were all purchased from Sigma-Aldrich (Missouri, United States). The calcium chelator Fluo-4-AM was purchased from Thermo Fisher Scientific (Massachusetts, United States). cGMP, PET-cGMP and 1-NH_2_-cGMP were purchased from BIOLOG Life Science Institute (Bremen, Germany).

Rat monoclonal anti-HA tag antibody (clone 3F10) was purchased from Roche LifeScience (Penzberg, Germany) and rabbit anti-human PKG antibody from Enzo life sciences (New York, United States). A rabbit polyclonal antibody against MSP1-30 (Woehlbier et al., 2006) as well as a rabbit anti-AMA1 antibody raised against the ectodomain (Collins et al., 2009) and rabbit anti-SERA5 (Stallmach et al., 2015) were all described previously. A mouse monoclonal antibody to Plasmepsin V was a kind gift from Daniel Goldberg (Washington University School of Medicine in St. Louis, Missouri, United States).

### *P. falciparum* culture and synchronisation

*P. falciparum* asexual blood stages were cultured in human erythrocytes (National Blood Transfusion Service, London, United Kingdom) and complete medium (CM) consisting of RPMI-1640 medium (Life Technologies, California, United States) supplemented with 0.5% AlbuMAX type II (Gibco), 50 μM hypoxanthine and 2 mM L-glutamine. Parasite cultures were incubated at 37°C and gassed with 90% N_2_, 5% CO_2_ and 5% O_2_ according to standard procedures (Trager and Jensen, 1976). Parasitemias were routinely monitored by examination of thin blood films fixed with 100% methanol and stained with 10% Giemsa stain in phosphate buffer (8 mM KH_2_PO_4_, 6 mM Na_2_HPO_4_, pH 7.0).

Tightly synchronous parasites were obtained by purifying segmented schizonts on a 70% isotonic Percoll (GE Healthcare, Illinois, United States) cushion and allowing them to invade fresh erythrocytes for 1-2 hours while shaking. Unruptured schizonts were lysed by treating with 5% D-sorbitol (Sigma, Missouri, United States) for 10 minutes (Lambros and Vanderberg, 1979) to obtain highly pure and synchronous ring stage cultures.

Induction of DiCre activity was achieved by treating early ring stage parasites (2-10 hours post invasion) with 50 nM RAP for 2-3 hours. Control parasites were treated with the equivalent volume of the vehicle DMSO (0.5% v/v).

### Transfection of *P. falciparum* schizonts

Highly synchronous late stage schizonts were used for transfection as previously described (Collins et al., 2013a) using the AmaxaTM 4D-Nucleofector system (Lonza, Basel, Switzerland). For each transfection ∼1.25×10^8^ Percoll-enriched schizonts were resuspended in supplemented P3 primary cell solution containing 20-50 μg of plasmid DNA and transferred to a Nucleocuvette. Parasites were electroporated using the FP158 setting then transferred back into culture. Modified parasites were selected 24 hours post transfection by addition of WR99210 (2.5 nM). WR99210 was removed seven days later when selecting for the presence Cas9/gRNA plasmids (GCα:HA:cKO and ATPase mutation) or left on until parasites emerged and then cycled on/off WR (GCα:HA).

### Plasmid construction

Primers used throughout this study were ordered from Integrated DNA Technologies (IDT, Iowa, United States) and are listed in Supplementary Table 1. To generate the GCα:HA parasite line, a 1.9 kb fragment corresponding to the 3’ end of the GCα coding region was PCR amplified with primers 14 and 15 and cloned into pHH1_PreDiCre_A_deltaH_deltaE (Collins et al., 2013a) via *Eco*RV and *Xho*I restriction sites, upstream of the sequence encoding a triple haemagglutinin (3xHA) tag. The resulting plasmid pHH1_PreDiCre_GCα-3xHA was transfected into the 3D7/1G5DiCre line constitutively expressing dimerisable Cre recombinase (Collins et al., 2013a). Transfected cultures were selected with WR99210 and then subjected to drug cycling to enrich for parasites having integrated the plasmid via single crossover recombination. Cultures were finally treated with RAP to activate Cre recombinase to recycle the hdhfr resistance marker. A clonal GCα:HA line sensitive to WR99210 was selected and further modified by CRISPR/Cas9-mediated gene editing to introduce a *loxPint* sequence into the ATPase domain of GCα to generate the GCα:HA:cKO line. A pUC19-based repair template was generated by first amplifying a 545 bp 5’ homology region and a 627 bp 3’ homology region from genomic DNA using primer pairs 16/17 and 18/13, respectively. The two PCR products were fused by overlap extension PCR using primers 16/13 and InFusion cloned into the *Hin*dIII and *Eco*RI sites of pUC19. A synthetic recodonised region of the GCα gene from bp 2109 to 2340 containing a SERA2-derived loxPint was ordered as a gBlock (IDT) and InFusion cloned into the *Afl*II and *Bam*HI sites located between the 5’ and 3’ homology regions. The repair template was linearised using *Pvu*I and transfected along with three pooled pDC2 plasmids each encoding the Cas9 protein, the hDHFR selection cassette, which confers resistance to the antifolate WR99210, and a unique sgRNA sequence targeting either TTTAATATGTGTTCTATAGC, TCTATAGCAGGAAAAACATA or CATATTCATCATAATCATTT. To mutate the ATPase domain of GCα, the repair template used to introduce the *loxPint* was modified to introduce either a D756N mutation, which would block formation of the aspartyl phosphate intermediate; or a wild type D756 synonymous mutation, which would serve as a control. The repair templates were generated by replacing the *loxPint* flanked by the *Bgl*II and *Kpn*I sites with overlap extension PCR products from primer sets 19/20 and 21/22 to introduce the D756N and D756 alleles, respectively. Each repair template plasmid was linearised with *Pvu*I, combined with the pool of three pDC2 Cas9 plasmids mentioned above, and transfected into wild type 3D7 parasites.

### Limiting dilution to generate clonal parasite lines

Clonal parasite lines were obtained by limiting dilution combined with a plaque formation readout as previously described (Thomas et al., 2016). The hematocrit and parasitemia of parasite cultures were determined by using a hemocytometer and by counting Giemsa-stained thin blood smears. Briefly, parasite cultures were diluted to give 0.3 parasites in 200 μl of culture at 1% hematocrit per well in a 96-well flat-bottom plate. After 9 days, plaque formation was assessed using an EVOS FL Cell Imaging System (Thermo Fisher Scientific). Wells containing single plaques were subsequently expanded and analysed by PCR.

### Diagnostic PCRs

Integration of the 3xHA-tagging construct into the GCα locus was confirmed using primers 1/2. RAP-induced excision of the hDHFR cassette to create the GCα:HA line was validated using primers 1/4. Integration of the artificial *loxPint* into the ATPase domain of the GCα locus to generate the GC:HA:cKO line was confirmed using primers 5/7, while primers 5/6 were used to detect the presence of wild type locus. Cre-mediated excision was validated using prime pairs 10/11 and 12/11 to detect the un-excised and excised loci, respectively. Primers 8/9, which amplify a segment in an unmodified distal locus, served as a DNA quality control.

### SYBR Green growth inhibition assays

To determine the effect of various test compounds on parasite growth, their half maximal effective concentration (EC_50_) values were determined by using the SYBR Green growth inhibition assay adapted from (Smilkstein et al., 2004). Test compounds were added as a series of 2-fold serial dilutions in triplicate to 96-well flat-bottom plates. Wells containing no drug or 10 nM chloroquine were also included in each plate and served as negative and positive controls, respectively. Synchronous ring stage parasites were added to achieve a starting parasitemia of 2% at 1% hematocrit and incubated at 37°C in a sealed gassed box for 72 hours. Parasites were then lysed in buffer containing 20 mM Tris, 5 mM EDTA, 0.008% saponin, 0.08% Triton X-100 and 1x SYBR Green I (Molecular Probes, Oregon, United States) at pH 7.5 and incubated for one hour at room temperature. SYBR Green fluorescence was measured using a Spectramax M3 plate reader (Molecular Devices, California, United States) with excitation and emission wavelengths of 485 and 535 nm, respectively. EC_50_ values were determined by non-linear regression analysis.

### FACS analysis to measure parasite growth and DNA content

Parasite cultures were seeded in triplicate wells per condition and samples were fixed in 4% formaldehye, 0.1% glutaraldehyde in PBS containing 1x SYBR Green I (Molecular Probes) and stored at 4°C overnight. The fixative was aspirated and the cells washed in PBS then analysed using a BD LSR II flow cytometer (BD Biosciences, New Jersey, United States), with 50,000 events collected for each sample. FlowJo 7 analysis software (FlowJo LLC, Oregon, United States) was used to analyse the data.

### Immunofluorescence microscopy

Air-dried thin blood smears were fixed in 4% formaldehyde in PBS for 20 minutes at room temperature followed by permeabilisation with 0.1% Triton X-100 in PBS. Blocking and antibody binding steps were performed in PBS containing 3% bovine serum albumin (BSA). Dual staining experiments were performed sequentially, starting with rat anti-HA, to eliminate cross-reactivity of the anti-rat secondary antibody with mouse or rabbit IgG. Secondary antibodies used were anti-rat IgG conjugated to Alexa Fluor 488, anti-rabbit IgG Alexa Fluor 594, and anti-mouse IgG Alexa Fluor 594, all highly cross-adsorbed. Slides were mounted in ProLong™ Gold Antifade Mountant containing 4’,6’-diamidino-2-phenylindole (DAPI; Thermo Fisher Scientific). Images were acquired at 100x magnification using a Nikon Eclipse Ti fluorescence microscope fitted with a Hamamatsu C11440 digital camera, and overlaid in ICY bioimage analysis software (icy.bioimageanalysis.org).

### Time-lapse video microscopy

Parasite egress was monitored by differential interference contrast (DIC) coupled with fluorescence microscopy using a Nikon Eclipse Ti fluorescence microscope with a 60x oil immersion objective and fitted with a Hamamatsu C11440 digital camera. Segmented schizonts treated with C2 (1.5 μM) overnight were Percoll enriched and resuspended in warm complete medium at 0.4% hematocrit, transferred to pre-warmed Poly-L-Lysine μ-Slide VI 0.4 (IBIDI, Planegg, Germany) imaging chambers and imaged on a temperature-controlled microscope stage held at 37°C. To visualise DMSO- and RAP-treated parasites simultaneously, one culture was stained with 1 µg/ml Hoechst 33342 prior to washing off C2 and pooling the cultures, as previously described (Das et al., 2015). Images were taken every 5 seconds for a total of 20-30 minutes, and the resulting videos were processed and analysed in ICY bioimage analysis software (icy.bioimageanalysis.org).

### Microscopy of Giemsa-stained blood films

Thin blood films fixed with 100% methanol and stained with 10% Giemsa stain in phosphate buffer (8 mM KH_2_PO_4_, 6 mM Na_2_HPO_4_, pH 7.0) were imaged using an Olympus BX51 microscope fitted with an Olympus SC30 digital colour camera through a 100× oil immersion objective. Images were processed in Graphic (Picta Inc.).

### Parasite protein extraction, SDS PAGE, and immunoblotting

Saponin-released parasites were lysed in 4 pellet volumes of CoIP buffer (150 mM NaCl, 0.5 mM EDTA, 1% NP40 and 10 mM Tris (pH 7.5)) supplemented with cOmplete EDTA-free protease inhibitor (Roche, Basel, Switzerland). Samples were incubated on ice for 10 minutes, centrifuged at 12,000 x g for 10 minutes at 4°C and the supernatant collected. Reducing SDS sample buffer was added and proteins resolved on 4%-15% Mini-PROTEAN TGX Stain-Free Precast Gels (Bio-Rad, California, United States) or 3%-8% NuPAGE Tris-Acetate Protein Gels (Thermo Fisher Scientific) for high molecular weight proteins. Proteins were transferred onto nitrocellulose membranes using a semidry Trans-Blot Turbo Transfer System (Bio-Rad) and blocked using 10% skimmed milk in PBS containing 0.1% Tween-20 (PBST). Antibody incubations were carried out in 1% skimmed milk in PBST and washed in PBST. Following incubation with secondary antibodies conjugated to near infrared (NIR) dyes, washed membranes were dried between Whatman 3MM blotting papers and imaged using an Azure c600 Imaging System (Azure Biosystems, California, United States) or a ChemiDoc Imaging System (Bio-Rad).

### Egress assays

Highly synchronous mature segmented schizonts from DMSO- and RAP-treated GCα:HA:cKO cultures that were treated with C2 (1.5 μM) at the early trophozoite stage were enriched on a Percoll gradient and washed several times in pre-warmed RPMI. Parasites were resuspended in RPMI at 3.25×10^8^ parasites/ml and 65 μl aliquots were dispensed in Eppendorf tubes. To harvest samples at each time point, parasites were pelleted at 9,000 x g and culture supernatants were purified using 0.22 μm Costar Spin-X centrifuge filters (Corning, New York, United States). The parasite pellets from the first time point were retained as a parasite loading control. Samples were subjected to western blot analysis, and probed for SERA5 as a measure of merozoite egress.

### Calcium release assays

Mature segmented schizonts from RAP- and DMSO-treated cultures were Percoll enriched and ∼1.25×10^8^ cells from each condition incubated in phenol red free RPMI containing 10 μM Fluo-4-AM (Invitrogen, California, United States) in the dark at 37°C for 45 minutes. The parasites were washed twice in pre-warmed phenol red free RPMI, then incubated for 20 minutes to allow for de-esterifcation of the AM ester. The parasites were washed twice and resuspended in phenol red free RPMI at 1.25×10^8^ parasites/ml. 100 μl of resuspended parasites we re-added to wells on the bottom half of a 96-well plate. Three wells containing phenol red free RPMI were also included as a control. Baseline Fluo-4 fluorescence in each well was read at 22 second intervals for 3 minutes using a Spectramax M3 plate reader (Molecular Devices) pre-warmed to 37°C with excitation and emission wavelengths of 483 and 525 nm, respectively. The plate was removed from the reader onto a heat block pre-warmed to 37°C and the parasites were resuspended and transferred to wells containing test compounds to give the desired final concentrations of zaprinast (75 μM), ionophore A23187 (20 μM) and DMSO (1.5%). The plate was placed back in the reader and read for a further 5 minutes at 22 second intervals. All samples were run in triplicate. Relative fluorescence units from reads at each time point and condition were averaged and baseline and DMSO control values subtracted.

### Measurement of intracellular cyclic nucleotide levels

Intracellular cyclic nucleotide levels in mature segmented schizonts were measured using ELISA-based high-sensitivity direct cAMP and cGMP colorimetric assay kits (Enzo life sciences). Around 1.25×10^8^ schizonts were Percoll-purified were obtained from RAP- and DMSO-treated cultures to which C2 (1.5 μM) had been added to prevent schizont rupture. The purified schizonts were incubated for 3 minutes in RPMI containing C2 only or C2 in the presence of the PDE inhibitor zaprinast (75 μM). Parasites were then pelleted at 9,000 x g, resuspended in 100 μl of 0.1 M HCl and incubated for 10 minutes at room temperature with intermittent vortexing to complete cell lysis. The samples were pelleted at 9,000 x g and the supernatant collected and stored at −80°C until required. Once all biological replicates were collected, each sample was diluted by adding 400 μl of 0.1 M HCl. Samples and standards were acetylated in order to improve sensitivity, according to the manufacturer’s instructions.

The detection ranges were 0.078 - 20 and 0.08 - 50 pmol/ml for the cAMP and cGMP assays, respectively. All samples and standards were set up in duplicate. Absorbance was measured at 405 nm using a Spectramax M3 plate reader (Molecular Devices).

### Treatment of parasite cultures with cGMP analogues to rescue GCα KO phenotype

Highly synchronous mature segmented schizonts from RAP-treated cultures at ∼48 h post invasion were treated with cGMP, 1-NH_2_-cGMP or PET-cGMP at concentrations ranging from 15.6 to 250 μM. Giemsa-stained thin blood smears were taken after 2 h and parasites were scored for their viability based on morphology. Parasites in 10 microscopic fields per condition were assigned to schizont, merozoite / pyknotic intracellular, or ring stage. Counts were performed blind by two researchers. To measure the effect of 30 μM PET cGMP on wild type and GCα KO replication rates, the cGMP analogue was added to synchronous DMSO and RAP-treated GCα:HA:cKO cultures when segmented schizonts appeared, washed off 10 h later and parasites harvested for FACS analysis at the ensuing early schizont stage.

### Sequence Alignments

Sequence alignments were performed using Clustal Omega, modified manually and guided by the alignment presented in a recent P4-ATPase structural study (Hiraizumi et al., 2019).

### Data analysis and statistical significance tests

GraphPad Prism 7 was used for all statistical analyses. The numbers of biological and technical replicates for each experiment are noted in the figure legends.

## Acknowledgments

We thank Oriana Kreutzfeld for a contribution to plasmid construct generation. We also thank Daniel Goldberg for kindly gifting the Plasmepsin V antibody.

---

**Supplementary Figure 1.**
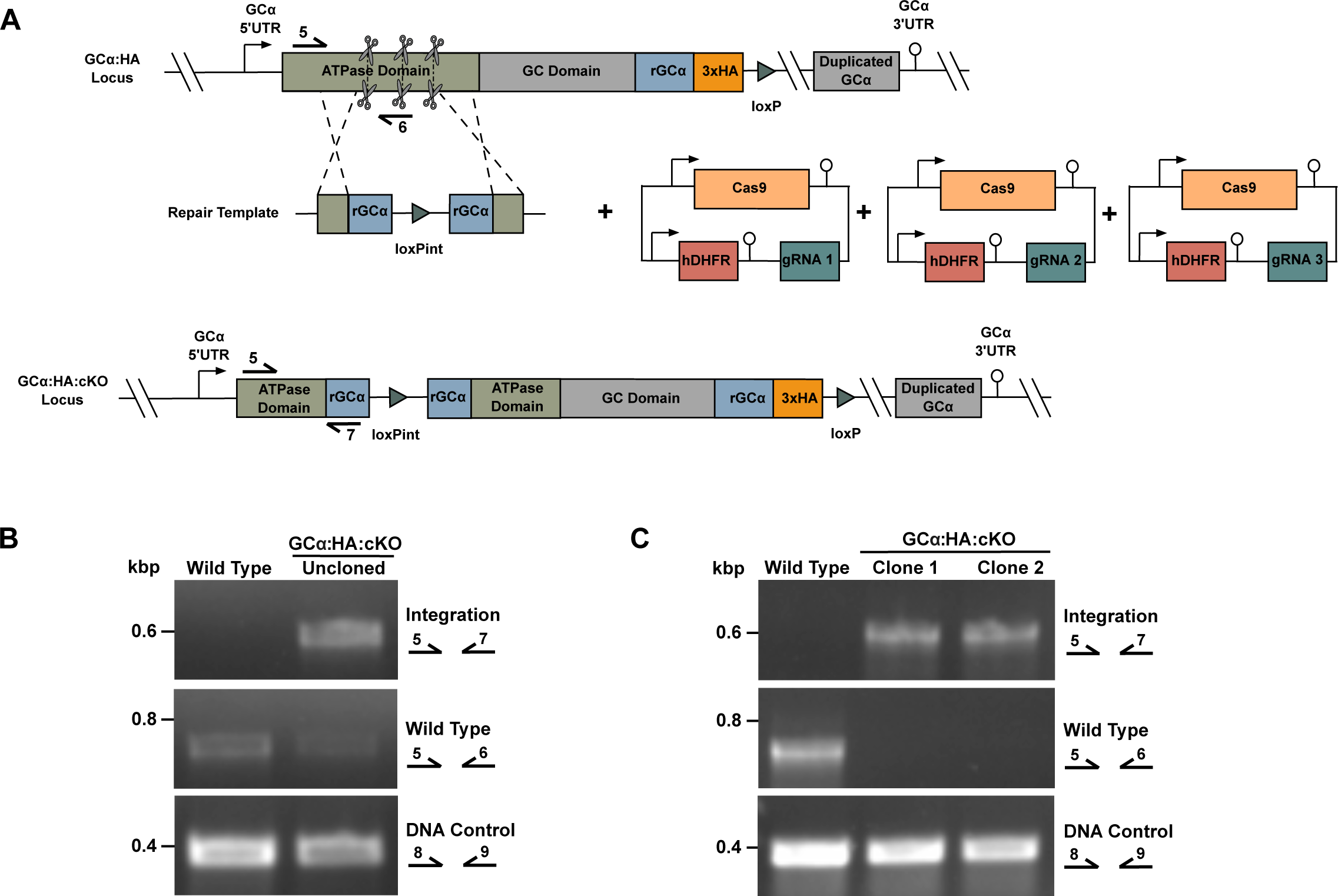
Generation of a GCα:HA:cKO line. **(A)** Schematic representation of the marker-free CRISPR/Cas9-mediated approach used to introduce the *loxPint* into the ATPase domain of *GC*α in the GCα:HA line to generate GCα:HA:cKO. Scissors indicate CRISPR/Cas9 cleavage sites, while arrows with numbers represent the relative position of oligonucleotide primers used for diagnostic PCR. A pool of three Cas9 plasmids harbouring different gRNA cassettes to target Cas9 to the ATPase domain of the *GCα* gene (pDC2-Cas9-hDHFR gRNA1-3) were co-transfected with a linearised repair template containing the *loxPint* sequence flanked by recodonised (*rGCα*) and endogenous *GCα* sequences, the latter serving as template for homology-directed repair. Transfected cultures were initially maintained in the presence of WR99210 to select for parasites harbouring one or more of the pDC2-Cas9-hDHFR plasmids. Promoters/5’UTRs and 3’UTRs/terminators are indicated by arrows and lollipops, respectively. **(B)** Diagnostic PCR analysis showing successful integration of the *loxPint* in the uncloned GCα:HA:cKO parasite population as well as the presence of GCα wild type locus. A DNA control PCR was included, which amplified a small segment from an independent locus (Pf3D7_1342600) to confirm the quality of the DNA used. PCR primers used are indicated on the right and their approximate binding sites within the modified GCα locus are shown in (A). **(C)** Diagnostic PCR analysis confirming successful integration of the *loxPint* and absence of wild type locus in two GCα:HA:cKO clones. A DNA control PCR was included, which amplified a small segment at an independent locus (Pf3D7_1342600) to confirm the quality of the DNA used. Approximate binding sites of specific PCR primers are shown in (A).

**Supplementary Figure 2.**
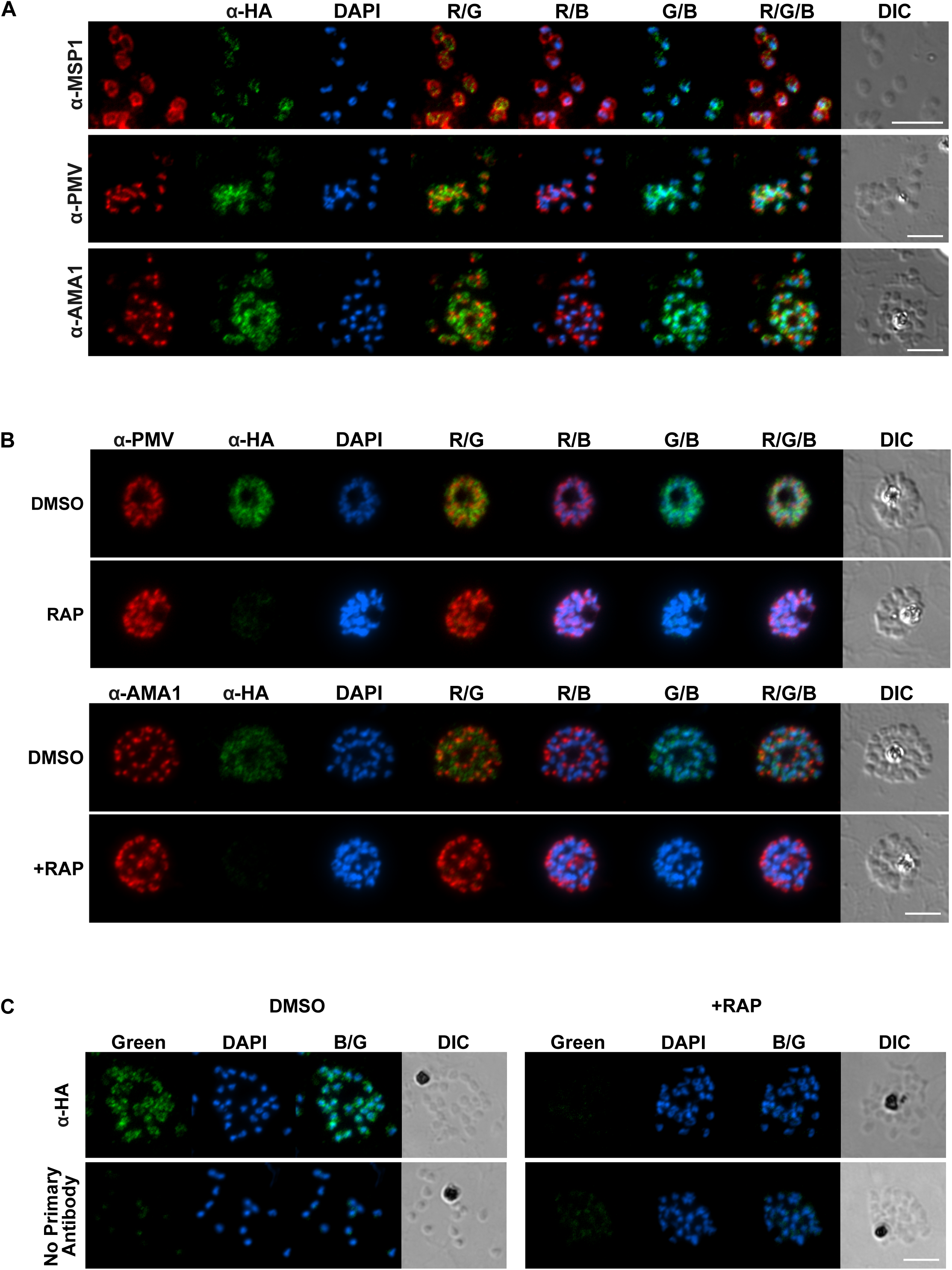
Subcellular localisation of GCα in schizonts and free merozoites, verification of GCα knockout at single cell level, and confirmation of specificity of the α -HA signal. **(A)** Dual staining IFA analysis of ruptured GCα:HA schizonts or free merozoites. Formaldehyde-fixed thin films were stained with α-HA (green) and co-stained with antibodies to markers for known subcellular compartments (red): MSP1 (parasite plasma membrane, top panel), plasmepsin V (endoplasmic reticulum, middle panel), and AMA1 (micronemes, bottom panel). Scale bar, 5 μm. **(B)** Dual staining IFA analysis of mature GCα:HA:cKO schizonts from cultures treated with either DMSO or RAP at ring stage of the same cycle. Slides were stained for GCα (α-HA, green) and co-stained for either plasmepsin V (upper panel) or AMA1 (lower panel), both red. Scale bar, 5 μm. **(C)** IFA analysis to verify specificity of α-HA signal observed for GCα-HA in schizonts. Slides containing GCα:HA:cKO schizonts from cultures either treated with DMSO or RAP were stained with α-HA followed by fluorescently labelled α-rat IgG (upper panels) or the same secondary antibody alone (lower panels). Scale bar, 5 μm.

**Supplementary Figure 3.**
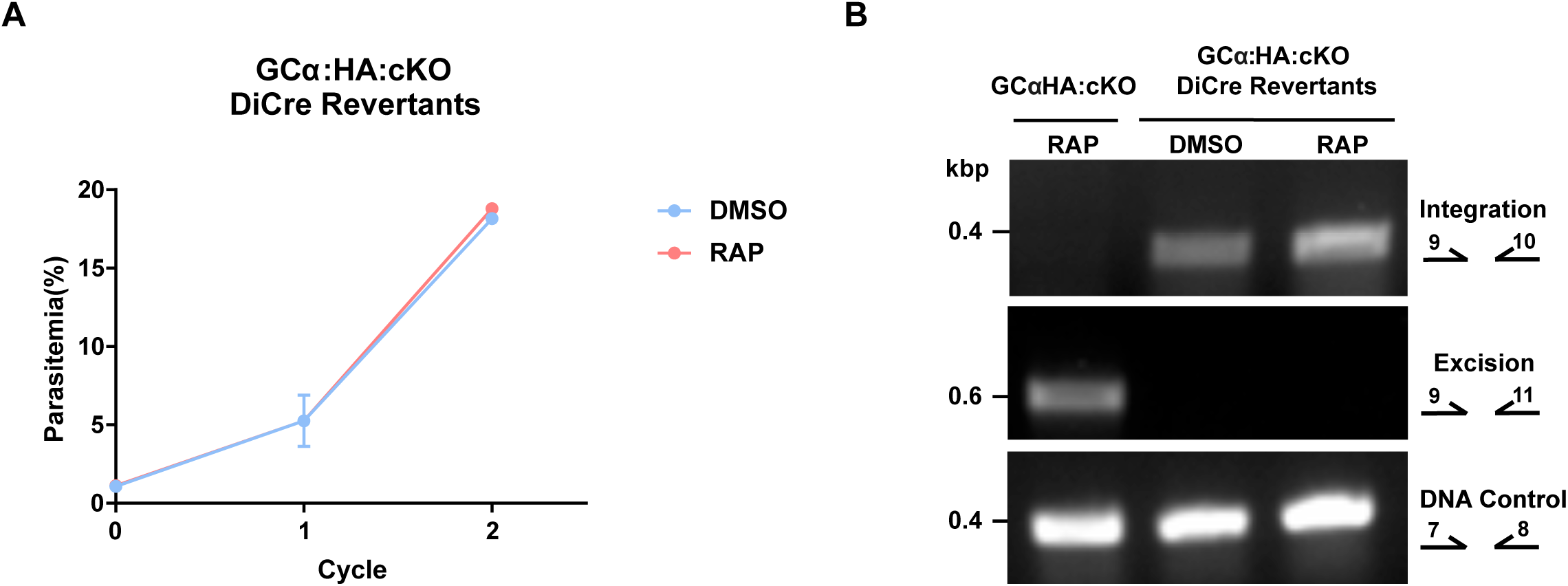
Parasites emerging from RAP-treated GCα:HA:cKO cultures are DiCre revertants with an intact GCα locus. **(A)** Around 16 days post RAP treatment, parasites emerged from GCα:HA:cKO cultures. Parasites recovered from these cultures were treated with either DMSO or RAP and growth measured over two cycles. Parasitaemias were determined by flow cytometry-based counting of SYBR Green positive cells. Data points plotted are means from two repeat experiments, each performed in triplicate. Error bars, S.D. RAP-treatment had no effect on parasite growth. **(B)** Genomic DNA collected at the end of a growth experiment as in (**A**) was analysed by diagnostic PCR along with a control obtained from the initial excision cycle (lane 1, RAP). No excised GCα locus was detected in parasites recovered from a previously RAP-treated culture (lane 2, DMSO) or after a second RAP treatment (lane 3, RAP). This strongly indicates that the recovered parasites represent DiCre revertants, a phenomenon previously observed on the 1G5 DiCre background. A DNA control PCR reaction was included, which amplified a small segment from an independent locus to test the quality of the DNA used.

**Supplementary Figure 4.**
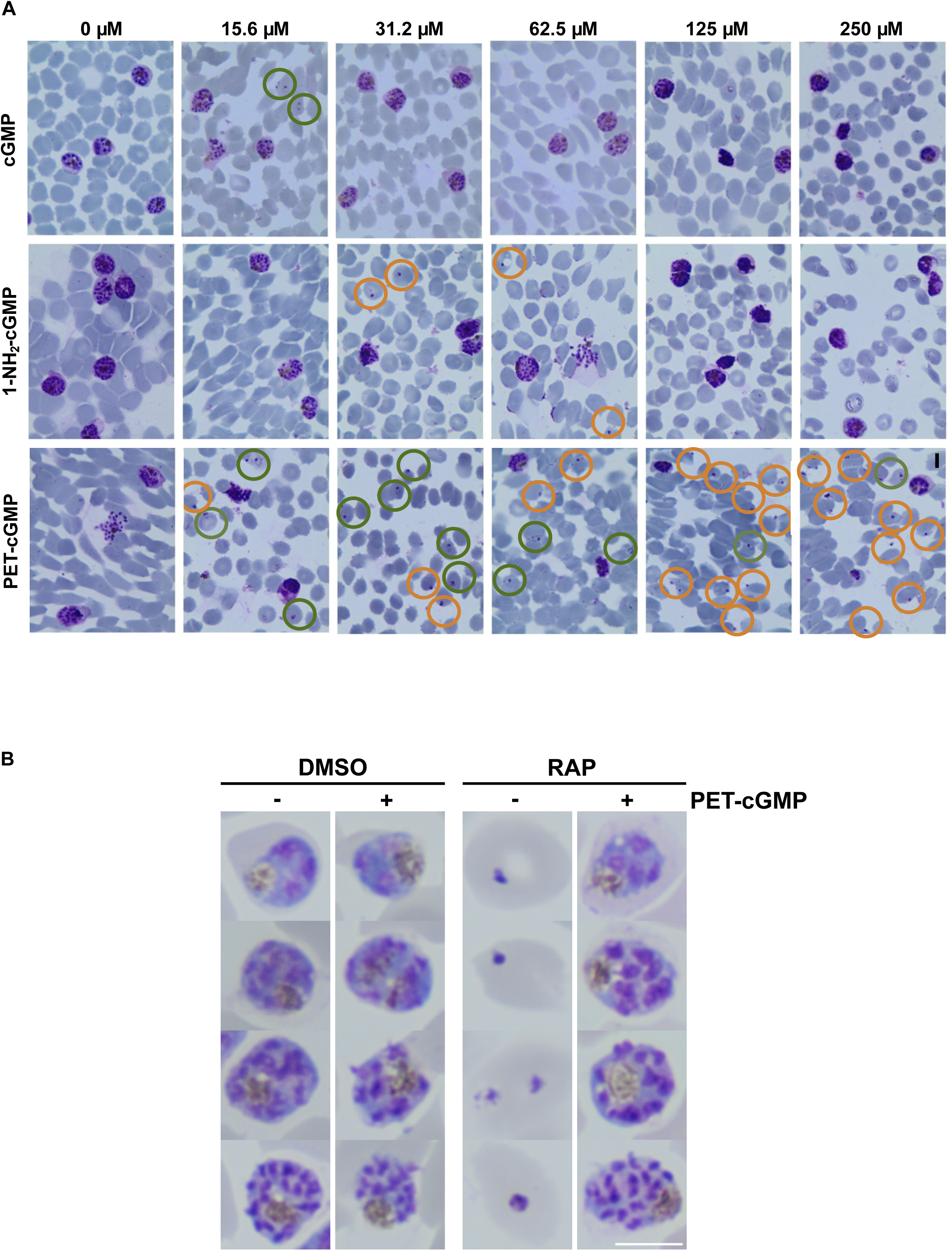
Rescue of GCα KO egress phenotype by addition of cGMP analogues. **(A)** Representative images of Giemsa-stained blood smears from RAP-treated GCα:HA:cKO cultures treated with various concentrations of cGMP, 1-NH_2_-cGMP and PET-cGMP at mature schizont stage. Samples were examined after 10 h. cGMP analogues added are indicated on the top and final concentrations to the left. Successful invasion events (newly formed ring stages) are circled in green, extracellular (merozoites) and pycnotic forms in orange. **(B)** Giemsa images of GCα:HA:cKO schizonts in the cycle post excision (cycle 1). Representative images are shown from DMSO and RAP-treated cultures with and without the addition of 30 μM PET-cGMP. PET-cGMP was added at the mature schizont stage of the excision cycle (cycle 0) and washed off 10 h later, when most parasites had egressed and re-invaded. Scale bar = 5 μm.

**Supplementary Figure 5.**
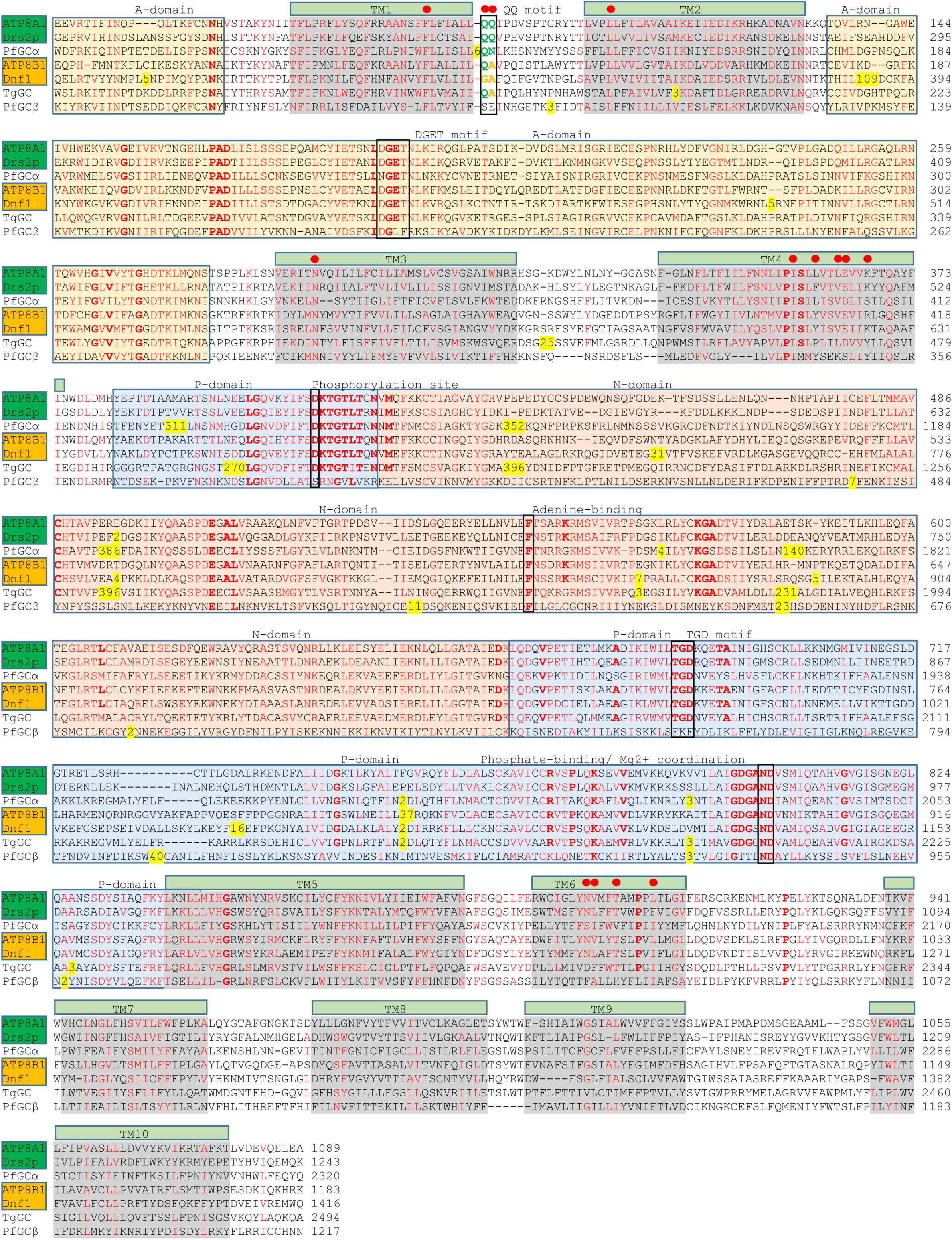
ATPase-like domain alignment. An alignment of the ATPase-like domain of *P. falciparum* GCα (PlasmoDB identifier: PF3D7_1138400) with human (ATP8A1, UniProt: Q9Y2Q0-2; ATP8B1, UniProt: O43520) and yeast (Drs2p, UniProt: P39524; Dnf1, UniProt: P32660) P4-ATPases to highlight functionally important residues. The ATPase-like domain of the *Toxoplasma gondii* GC (accession number EPT31724) and that of *P. falciparum* GCβ (PlasmoDB identifier: PF3D7_1360500) are also included in the alignment for comparison. The names of yeast/mammalian P4-ATPases that flip PS are coloured in green and those that flip PC are coloured in amber. Amino acids associated with selectivity for PS or PC flipping in human and yeast PS-ATPases are coloured in green and amber respectively. The positions of amino acids shown to be important for phospholipid binding/translocation in a crystal structure of human ATP8A1 are indicated with red circles above the sequence (mostly conserved in GCα). The important ATP-binding amino acids of the ATP8A sequence (which are all conserved in GCα) are F534 in the N-domain that contacts the adenine ring; the phosphate group interacts with the D409 and T411 in the DKTG motif containing the phosphorylation site (D409) and the N789 and the D790 towards the end of the P domain along with a Mg^2+^ ion. The strong association of the A domain with the phosphorylation site in the human ATP8A crystal structure is mediated by D189 and G190 in the DGET motif where the aspartate is replaced by an asparagine in GCα. Following the scheme depicted in the structural study of human ATP8A1, the actuator domain (A-domain) sequence is shown in an orange box, phosphorylation domain (P-domain) sequence in a blue box and the nucleotide-binding domain (N-domain) sequence in a pink box. Important functional motifs (described in the Results section) are indicated above the sequence and boxed. Transmembrane domains are indicated above the sequence by green bars and sequences are highlighted in grey. Amino acids that are identical in the four human and yeast P4-ATPases as well as P2-ATPases (SERCA Ca^2+^ ATPase, UniProt: P04191 and Na^+^/K^+^ ATPase, UniProt: Q4H132, not included in the alignment) are coloured in bold red. Residues identical or similar in 5 of the 7 aligned sequences are coloured red to show highly conserved regions. Amino acid residue are shown to the right of each sequence.

**Supplementary Table 1.**
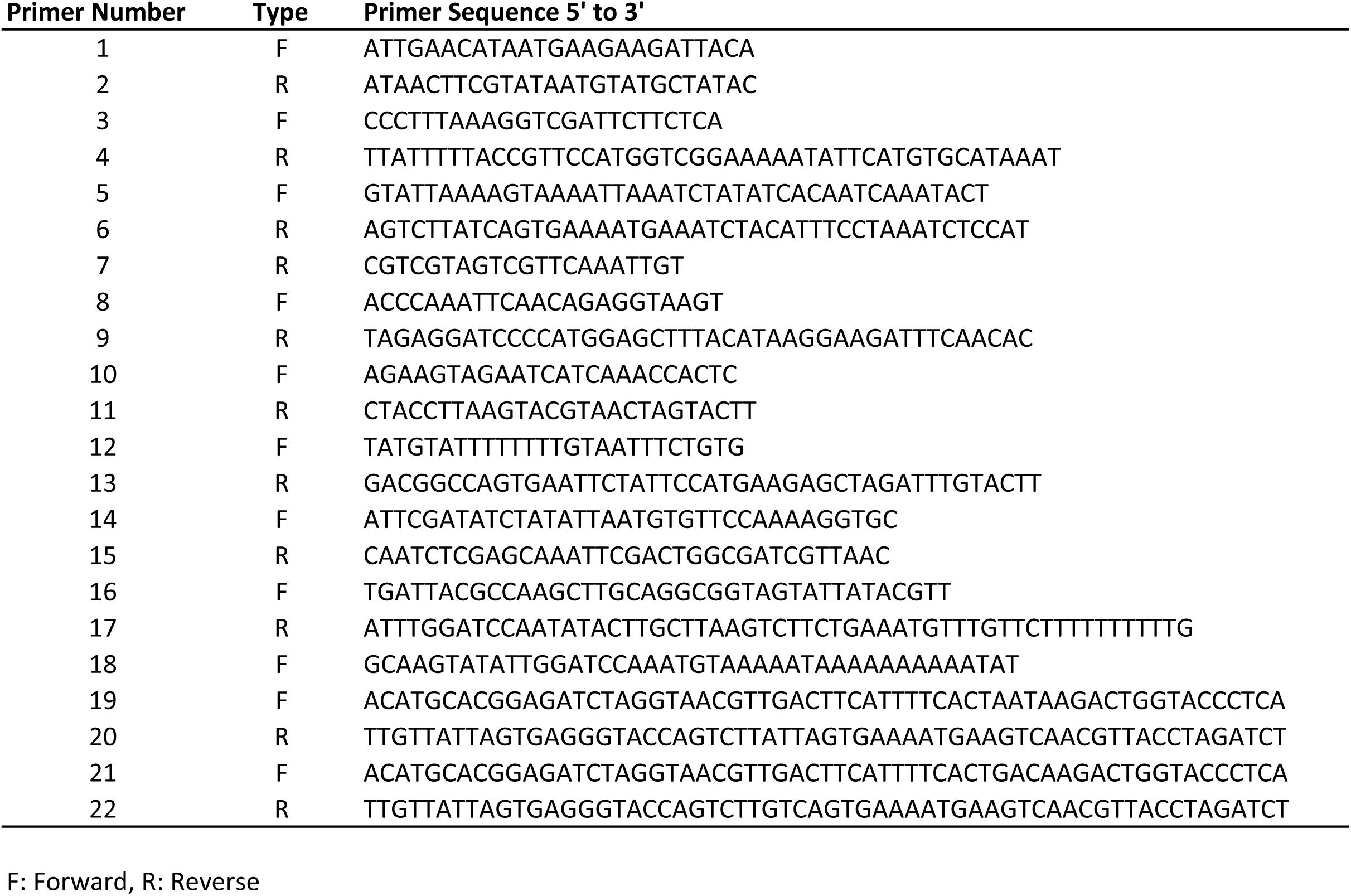

## References

Alam, M. M., Solyakov, L., Bottrill, A. R., Flueck, C., Siddiqui, F. A., Singh, S., Mistry, S., Viskaduraki, M., Lee, K., Hopp, C. S., Chitnis, C. E., Doerig, C., Moon, R. W., Green, J. L., Holder, A. A., Baker, D. A. & Tobin, A. B. 2015. Phosphoproteomics reveals malaria parasite Protein Kinase G as a signalling hub regulating egress and invasion. Nat Commun, 6, 7285.

Baker, D. A., Drought, L. G., Flueck, C., Nofal, S. D., Patel, A., Penzo, M. & Walker, E. M. 2017. Cyclic nucleotide signalling in malaria parasites. Open Biol, 7.

Baker, D. A. & Kelly, J. M. 2004. Structure, function and evolution of microbial adenylyl and guanylyl cyclases. Mol Microbiol, 52, 1229–42.

Best, J. T., Xu, P. & Graham, T. R. 2019. Phospholipid flippases in membrane remodeling and transport carrier biogenesis. Curr Opin Cell Biol, 59, 8–15.

Billker, O., Lindo, V., Panico, M., Etienne, A. E., Paxton, T., Dell, A., Rogers, M., Sinden, R. E. & Morris, H. R. 1998. Identification of xanthurenic acid as the putative inducer of malaria development in the mosquito. Nature, 392, 289–92.

Bisio, H., Lunghi, M., Brochet, M. & Soldati-Favre, D. 2019. Phosphatidic acid governs natural egress in Toxoplasma gondii via a guanylate cyclase receptor platform. Nat Microbiol, 4, 420–428.

Brochet, M., Collins, M. O., Smith, T. K., Thompson, E., Sebastian, S., Volkmann, K., Schwach, F., Chappell, L., Gomes, A. R., Berriman, M., Rayner, J. C., Baker, D. A., Choudhary, J. & Billker, O. 2014. Phosphoinositide metabolism links cGMP-dependent protein kinase G to essential Ca(2)(+) signals at key decision points in the life cycle of malaria parasites. PLoS Biol, 12, e1001806.

Brown, K. M. & Sibley, L. D. 2018. Essential cGMP Signaling in Toxoplasma Is Initiated by a Hybrid P-Type ATPase-Guanylate Cyclase. Cell Host Microbe, 24, 804–816 e6.

Bushell, E., Gomes, A. R., Sanderson, T., Anar, B., Girling, G., Herd, C., Metcalf, T., Modrzynska, K., Schwach, F., Martin, R. E., Mather, M. W., Mcfadden, G. I., Parts, L., Rutledge, G. G., Vaidya, A. B., Wengelnik, K., Rayner, J. C. & Billker, O. 2017. Functional Profiling of a Plasmodium Genome Reveals an Abundance of Essential Genes. Cell, 170, 260–272 e8.

Byun, J. A., Van, K., Huang, J., Henning, P., Franz, E., Akimoto, M., Herberg, F. W., Kim, C. & Melacini, G. 2020. Mechanism of allosteric inhibition in the Plasmodium falciparum cGMP-dependent protein kinase. J Biol Chem.

Carucci, D. J., Witney, A. A., Muhia, D. K., Warhurst, D. C., Schaap, P., Meima, M., Li, J. L., Taylor, M. C., Kelly, J. M. & Baker, D. A. 2000. Guanylyl cyclase activity associated with putative bifunctional integral membrane proteins in Plasmodium falciparum. J Biol Chem, 275, 22147–56.

Collins, C. R., Das, S., Wong, E. H., Andenmatten, N., Stallmach, R., Hackett, F., Herman, J. P., Muller, S., Meissner, M. & Blackman, M. J. 2013a. Robust inducible Cre recombinase activity in the human malaria parasite Plasmodium falciparum enables efficient gene deletion within a single asexual erythrocytic growth cycle. Mol Microbiol, 88, 687–701.

Collins, C. R., Hackett, F., Strath, M., Penzo, M., Withers-Martinez, C., Baker, D. A. & Blackman, M. J. 2013b. Malaria parasite cGMP-dependent protein kinase regulates blood stage merozoite secretory organelle discharge and egress. PLoS Pathog, 9, e1003344.

Collins, C. R., Withers-Martinez, C., Hackett, F. & Blackman, M. J. 2009. An inhibitory antibody blocks interactions between components of the malarial invasion machinery. PLoS Pathog, 5, e1000273.

Das, S., Hertrich, N., Perrin, A. J., Withers-Martinez, C., Collins, C. R., Jones, M. L., Watermeyer, J. M., Fobes, E. T., Martin, S. R., Saibil, H. R., Wright, G. J., Treeck, M., Epp, C. & Blackman, M. J. 2015. Processing of Plasmodium falciparum Merozoite Surface Protein MSP1 Activates a Spectrin-Binding Function Enabling Parasite Egress from RBCs. Cell Host Microbe, 18, 433–44.

Falae, A., Combe, A., Amaladoss, A., Carvalho, T., Menard, R. & Bhanot, P. 2010. Role of Plasmodium berghei cGMP-dependent protein kinase in late liver stage development. J Biol Chem, 285, 3282–8.

Gao, H., Yang, Z., Wang, X., Qian, P., Hong, R., Chen, X., Su, X. Z., Cui, H. & Yuan, J. 2018. ISP1-Anchored Polarization of GCbeta/CDC50A Complex Initiates Malaria Ookinete Gliding Motility. Curr Biol, 28, 2763–2776 e6.

Gardner, M. J., Hall, N., Fung, E., White, O., Berriman, M., Hyman, R. W., Carlton, J. M., Pain, A., Nelson, K. E., Bowman, S., Paulsen, I. T., James, K., Eisen, J. A., Rutherford, K., Salzberg, S. L., Craig, A., Kyes, S., Chan, M. S., Nene, V., Shallom, S. J., Suh, B., Peterson, J., Angiuoli, S., Pertea, M., Allen, J., Selengut, J., Haft, D., Mather, M. W., Vaidya, A. B., Martin, D. M., Fairlamb, A. H., Fraunholz, M. J., Roos, D. S., Ralph, S. A., Mcfadden, G. I., Cummings, L. M., Subramanian, G. M., Mungall, C., Venter, J. C., Carucci, D. J., Hoffman, S. L., Newbold, C., Davis, R. W., Fraser, C. M. & Barrell, B. 2002. Genome sequence of the human malaria parasite Plasmodium falciparum. Nature, 419, 498–511.

Govindasamy, K., Jebiwott, S., Jaijyan, D. K., Davidow, A., Ojo, K. K., Van Voorhis, W. C., Brochet, M., Billker, O. & Bhanot, P. 2016. Invasion of hepatocytes by Plasmodium sporozoites requires cGMP-dependent protein kinase and calcium dependent protein kinase 4. Mol Microbiol, 102, 349–363.

Gunay-Esiyok, O., Scheib, U., Noll, M. & Gupta, N. 2019. An unusual and vital protein with guanylate cyclase and P4-ATPase domains in a pathogenic protist. Life Sci Alliance, 2.

Hirai, M., Arai, M., Kawai, S. & Matsuoka, H. 2006. PbGCbeta is essential for Plasmodium ookinete motility to invade midgut cell and for successful completion of parasite life cycle in mosquitoes. J Biochem, 140, 747–57.

Hiraizumi, M., Yamashita, K., Nishizawa, T. & Nureki, O. 2019. Cryo-EM structures capture the transport cycle of the P4-ATPase flippase. Science, 365, 1149–1155.

Hopp, C. S., Flueck, C., Solyakov, L., Tobin, A. & Baker, D. A. 2012. Spatiotemporal and functional characterisation of the Plasmodium falciparum cGMP-dependent protein kinase. PLoS One, 7, e48206.

Jia, Y., Marq, J. B., Bisio, H., Jacot, D., Mueller, C., Yu, L., Choudhary, J., Brochet, M. & Soldati-Favre, D. 2017. Crosstalk between PKA and PKG controls pH-dependent host cell egress of Toxoplasma gondii. EMBO J, 36, 3250–3267.

Jiang, Y., Wei, J., Cui, H., Liu, C., Zhi, Y., Jiang, Z., Li, Z., Li, S., Yang, Z., Wang, X., Qian, P., Zhang, C., Zhong, C., Su, X. Z. & Yuan, J. 2020. An intracellular membrane protein GEP1 regulates xanthurenic acid induced gametogenesis of malaria parasites. Nat Commun, 11, 1764.

Jones, M. L., Das, S., Belda, H., Collins, C. R., Blackman, M. J. & Treeck, M. 2016. A versatile strategy for rapid conditional genome engineering using loxP sites in a small synthetic intron in Plasmodium falciparum. Sci Rep, 6, 21800.

Kenthirapalan, S., Waters, A. P., Matuschewski, K. & Kooij, T. W. 2016. Functional profiles of orphan membrane transporters in the life cycle of the malaria parasite. Nat Commun, 7, 10519.

Koussis, K., Withers-Martinez, C., Baker, D. A. & Blackman, M. J. 2020. Simultaneous multiple allelic replacement in the malaria parasite enables dissection of PKG function. Life Sci Alliance, 3.

Kuhlbrandt, W. 2004. Biology, structure and mechanism of P-type ATPases. Nat Rev Mol Cell Biol, 5, 282–95.

Lakshmanan, V., Fishbaugher, M. E., Morrison, B., Baldwin, M., Macarulay, M., Vaughan, A. M., Mikolajczak, S. A. & Kappe, S. H. 2015. Cyclic GMP balance is critical for malaria parasite transmission from the mosquito to the mammalian host. MBio, 6, e02330.

Lambros, C. & Vanderberg, J. P. 1979. Synchronization of Plasmodium falciparum erythrocytic stages in culture. J Parasitol, 65, 418–20.

Linder, J. U., Engel, P., Reimer, A., Kruger, T., Plattner, H., Schultz, A. & Schultz, J. E. 1999. Guanylyl cyclases with the topology of mammalian adenylyl cyclases and an N-terminal P-type ATPase-like domain in Paramecium, Tetrahymena and Plasmodium. EMBO J, 18, 4222–32.

Lopez-Marques, R. L., Holthuis, J. C. & Pomorski, T. G. 2011. Pumping lipids with P4-ATPases. Biol Chem, 392, 67–76.

Lutsenko, S. & Kaplan, J. H. 1995. Organization of P-type ATPases: significance of structural diversity. Biochemistry, 34, 15607–13.

Mcrobert, L., Taylor, C. J., Deng, W., Fivelman, Q. L., Cummings, R. M., Polley, S. D., Billker, O. & Baker, D. A. 2008. Gametogenesis in malaria parasites is mediated by the cGMP-dependent protein kinase. PLoS Biol, 6, e139.

Michalakis, S., Becirovic, E. & Biel, M. 2018. Retinal Cyclic Nucleotide-Gated Channels: From Pathophysiology to Therapy. Int J Mol Sci, 19.

Montigny, C., Lyons, J., Champeil, P., Nissen, P. & Lenoir, G. 2016. On the molecular mechanism of flippase- and scramblase-mediated phospholipid transport. Biochim Biophys Acta, 1861, 767–783.

Moon, R. W., Taylor, C. J., Bex, C., Schepers, R., Goulding, D., Janse, C. J., Waters, A. P., Baker, D. A. & Billker, O. 2009. A cyclic GMP signalling module that regulates gliding motility in a malaria parasite. PLoS Pathog, 5, e1000599.

Moudy, R., Manning, T. J. & Beckers, C. J. 2001. The loss of cytoplasmic potassium upon host cell breakdown triggers egress of Toxoplasma gondii. J Biol Chem, 276, 41492–501.

Muhia, D. K., Swales, C. A., Deng, W., Kelly, J. M. & Baker, D. A. 2001. The gametocyte-activating factor xanthurenic acid stimulates an increase in membrane-associated guanylyl cyclase activity in the human malaria parasite Plasmodium falciparum. Mol Microbiol, 42, 553–60.

Park, M., Sandner, P. & Krieg, T. 2018. cGMP at the centre of attention: emerging strategies for activating the cardioprotective PKG pathway. Basic Res Cardiol, 113, 24.

Paul, A. S., Miliu, A., Paulo, J. A., Goldberg, J. M., Bonilla, A. M., Berry, L., Seveno, M., Braun-Breton, C., Kosber, A. L., Elsworth, B., Arriola, J. S. N., Lebrun, M., Gygi, S. P., Lamarque, M. H. & Duraisingh, M. T. 2020. Co-option of Plasmodium falciparum PP1 for egress from host erythrocytes. Nat Commun, 11, 3532.

Poulsen, L. R., Lopez-Marques, R. L. & Palmgren, M. G. 2008. Flippases: still more questions than answers. Cell Mol Life Sci, 65, 3119–25.

Roiko, M. S., Svezhova, N. & Carruthers, V. B. 2014. Acidification Activates Toxoplasma gondii Motility and Egress by Enhancing Protein Secretion and Cytolytic Activity. PLoS Pathog, 10, e1004488.

Roland, B. P. & Graham, T. R. 2016. Decoding P4-ATPase substrate interactions. Crit Rev Biochem Mol Biol, 51, 513–527.

Salowe, S. P., Wiltsie, J., Liberator, P. A. & Donald, R. G. 2002. The role of a parasite-specific allosteric site in the distinctive activation behavior of Eimeria tenella cGMP-dependent protein kinase. Biochemistry, 41, 4385–91.

Smilkstein, M., Sriwilaijaroen, N., Kelly, J. X., Wilairat, P. & Riscoe, M. 2004. Simple and inexpensive fluorescence-based technique for high-throughput antimalarial drug screening. Antimicrob Agents Chemother, 48, 1803–6.

Stallmach, R., Kavishwar, M., Withers-Martinez, C., Hackett, F., Collins, C. R., Howell, S. A., Yeoh, S., Knuepfer, E., Atid, A. J., Holder, A. A. & Blackman, M. J. 2015. Plasmodium falciparum SERA5 plays a non-enzymatic role in the malarial asexual blood-stage lifecycle. Mol Microbiol, 96, 368–87.

Taylor, C. J., Mcrobert, L. & Baker, D. A. 2008. Disruption of a Plasmodium falciparum cyclic nucleotide phosphodiesterase gene causes aberrant gametogenesis. Mol Microbiol, 69, 110–8.

Taylor, H. M., Mcrobert, L., Grainger, M., Sicard, A., Dluzewski, A. R., Hopp, C. S., Holder, A. A. & Baker, D. A. 2010. The malaria parasite cyclic GMP-dependent protein kinase plays a central role in blood-stage schizogony. Eukaryot Cell, 9, 37–45.

Thomas, J. A., Collins, C. R., Das, S., Hackett, F., Graindorge, A., Bell, D., Deu, E. & Blackman, M. J. 2016. Development and Application of a Simple Plaque Assay for the Human Malaria Parasite Plasmodium falciparum. PLoS One, 11, e0157873.

Thomas, J. A., Tan, M. S. Y., Bisson, C., Borg, A., Umrekar, T. R., Hackett, F., Hale, V. L., Vizcay-Barrena, G., Fleck, R. A., Snijders, A. P., Saibil, H. R. & Blackman, M. J. 2018. A protease cascade regulates release of the human malaria parasite Plasmodium falciparum from host red blood cells. Nat Microbiol, 3, 447–455.

Timcenko, M., Lyons, J. A., Januliene, D., Ulstrup, J. J., Dieudonne, T., Montigny, C., Ash, M. R., Karlsen, J. L., Boesen, T., Kuhlbrandt, W., Lenoir, G., Moeller, A. & Nissen, P. 2019. Structure and autoregulation of a P4-ATPase lipid flippase. Nature, 571, 366–370.

Trager, W. & Jensen, J. B. 1976. Human malaria parasites in continuous culture. Science, 193, 673–5.

Woehlbier, U., Epp, C., Kauth, C. W., Lutz, R., Long, C. A., Coulibaly, B., Kouyate, B., Arevalo-Herrera, M., Herrera, S. & Bujard, H. 2006. Analysis of antibodies directed against merozoite surface protein 1 of the human malaria parasite Plasmodium falciparum. Infect Immun, 74, 1313–22.

Yang, L., Uboldi, A. D., Seizova, S., Wilde, M. L., Coffey, M. J., Katris, N. J., Yamaryo-Botte, Y., Kocan, M., Bathgate, R. A. D., Stewart, R. J., Mcconville, M. J., Thompson, P. E., Botte, C. Y. & Tonkin, C. J. 2019. An apically located hybrid guanylate cyclase-ATPase is critical for the initiation of Ca(2+) signaling and motility in Toxoplasma gondii. J Biol Chem, 294, 8959–8972.

Zhang, M., Wang, C., Otto, T. D., Oberstaller, J., Liao, X., Adapa, S. R., Udenze, K., Bronner, I. F., Casandra, D., Mayho, M., Brown, J., Li, S., Swanson, J., Rayner, J. C., Jiang, R. H. Y. & Adams, J. H. 2018. Uncovering the essential genes of the human malaria parasite Plasmodium falciparum by saturation mutagenesis. Science, 360.

